# Cellular Stress Signaling Activates Type-I IFN Response Through FOXO3-regulated Lamin Posttranslational Modification

**DOI:** 10.1101/2020.05.03.075143

**Authors:** Inah Hwang, Ziwei Dai, Fei Li, Teresa Sanchez, Jason W Locasale, Lewis L Cantley, Hongwu Zheng, Jihye Paik

## Abstract

Neural stem/progenitor cells (NSPCs) persist over the lifespan while encountering constant challenges from age or injury related brain environmental changes, including elevated oxidative stress. A time-dependent stress response that regulates the dynamic balance between quiescence and differentiation is thus essential to preserve NSPC long-term regenerative potential. Here we report that acutely elevated cellular oxidative stress in NSPCs suppresses neurogenic differentiation through induction of FOXO3-mediated cGAS/STING and type I interferon (IFN-I) responses. We show that oxidative stress activates FOXO3 promoting upregulation of its transcriptional target glycine-N-methyltransferase (GNMT) and thus depletion of s-adenosylmethionine (SAM), a key co-substrate involved in methyl group transfer reactions. Mechanistically, we demonstrate that reduced intracellular SAM availability disrupts carboxymethylation and maturation of nuclear lamin, which trigger cytosolic release of chromatin fragments and subsequent activation of the cGAS/STING/IFN-I cascade. Together, our findings suggest the FOXO3-GNMT/SAM-lamin-cGAS/STING-IFN-I signaling cascade as a critical stress response program that preserves its long-term regenerative potential.

## Main

Stem cells persist over the total mammalian lifespan to maintain tissue homeostasis by replacing damaged or lost cells and the deterioration of their differentiation capacity is one of the key components of organismal aging^1^. Elevation of stress response pathways is a hallmark of aging tissues which also promotes the depletion of adult stem cells by inducing senescence or cell death^2–4^. Intrinsic and extrinsic cell-stressors such as DNA damage, mitochondrial dysfunction, loss of proteostasis, and the inflammatory tissue milieu contribute to an increased stress response. In particular, oxidative stress contributes to the functional decline of stem cells by inflicting damage to cellular macromolecules ultimately leading to cytostasis or cytotoxicity^1^. Model organism for precocious stem-cell depletion or dysfunction emphasize the role of key molecules involved in oxidative stress response (i.e. *Atm^5^, Foxo3^6–10^, Prdm16^11^*) in maintaining stem-cell reserves. Nevertheless, the direct cellular consequence of stress response that translates as molecular aging of stem cells remains broad and non-specific.

Among the many organs, the brain is particularly vulnerable due to its high oxygen consumption, unusual enrichment of polyunsaturated fatty acids as well as the presence of excitotoxic amino acids^12^. As a result, neural stem/progenitor-cells (NSPCs) constantly face stressful challenges and decline in their neurogenic potential in adult brains^13, 14^. However, the underlying mechanism by which elevated stress response regulates NSPC fate remains poorly understood.

FOXO transcription factors play evolutionarily conserved roles in a wide range of biological processes from aging to metabolism, not only by sensing stress but also through promoting stress resistance^15^. For example, previous studies indicated that FOXO is required for long-term regenerative potential of the hematopoietic stem cell (HSC) by regulating the response to physiologic oxidative stress and quiescence^7^. In the central nervous system, FOXO expression not only serves a key role in preserving neural stem cell pools^9, 10^ but also protects neurons against age-related axonal degeneration across species^16–18^. But despite these advances, there still lacks a mechanistic understanding of how aging-related oxidative stress affects FOXO activation systematically and whether and how that contributes to the neuroprotective responses.

The type-I interferon (IFN-I) response is an innate immune response that can be induced by a number of pattern recognition receptors^19^. Among them, cytosolic DNA fragments are recognized by cyclic GMP-AMP synthetase (cGAS) which initiates reaction of GTP and ATP to form cyclic GMP-AMP (cGAMP) cGAMP, a ligand of the signaling adaptor stimulator of interferon genes (STING, TMEM173). The binding of cGMP to STING activates TANK Binding Kinase 1 (TBK1) kinases-mediated phosphorylation of transcription factor IRF3 that triggers IFNα/β production and subsequent IFN response^20–23^. Increased IFN-I response has been shown to promote NSPC quiescence and suppress neurogenic differentiation^13, 24^. Interestingly, a recent study revealed that IFN-I signaling is elevated in the brain of aged humans and animals and correlates with increased oxidative stress^13^. But the connection between oxidative stress and IFN-I response is unclear.

Here, we report that oxidative stress-induced FOXO3 activation promotes transcriptional upregulation of N-methyltransferase GNMT to deplete S-adenosyl-L-methionine (SAM). Using the NSPC system, we further uncovered that reduction of intracellular SAM availability disrupts nuclear lamin maturation that eventually leads to cytosolic DNA leakage, cGAS/STING activation and IFN-I response. As a result, NSPC enters quiescence at the expense of neurogenic differentiation. These findings established FOXO3-GNMT/SAM-lamin-cGAS/STING-IFN-I signaling cascade as a critical stress response program that protects NSPC from detrimental environmental insults.

## Results

### High redox potential-mediated cellular stress activates IFN-I pathway

The neurogenic differentiation potential of NSPCs declines under iatrogenic insults, traumatic injuries, or inflammatory stress conditions^13, 25^. Among these, to determine how oxidative stress signal impacts NSPC differentiation, we subjected the cultured murine NSPCs to either pro-oxidant agent paraquat (PQ) or anti-oxidant N-acetylcysteine (NAC). Measurement by a ratiometric Grx-roGFP2 sensor confirmed that PQ treatment induced a marked elevation of intracellular redox potential relative to the mock-treated NSPCs, whereas NAC treatment led to a significant reduction (Fig. 1a, b). Importantly, compared to the mock-treated control NSPCs, we found that NSPCs under PQ but not NAC treatment, exhibited marked reduction in production of TUBB3 or doublecortin (DCX)-positive newly born neurons when induced to differentiate (Fig. 1c), suggesting a regulatory role of oxidative stress response on neurogenic differentiation.

**Figure 1.**
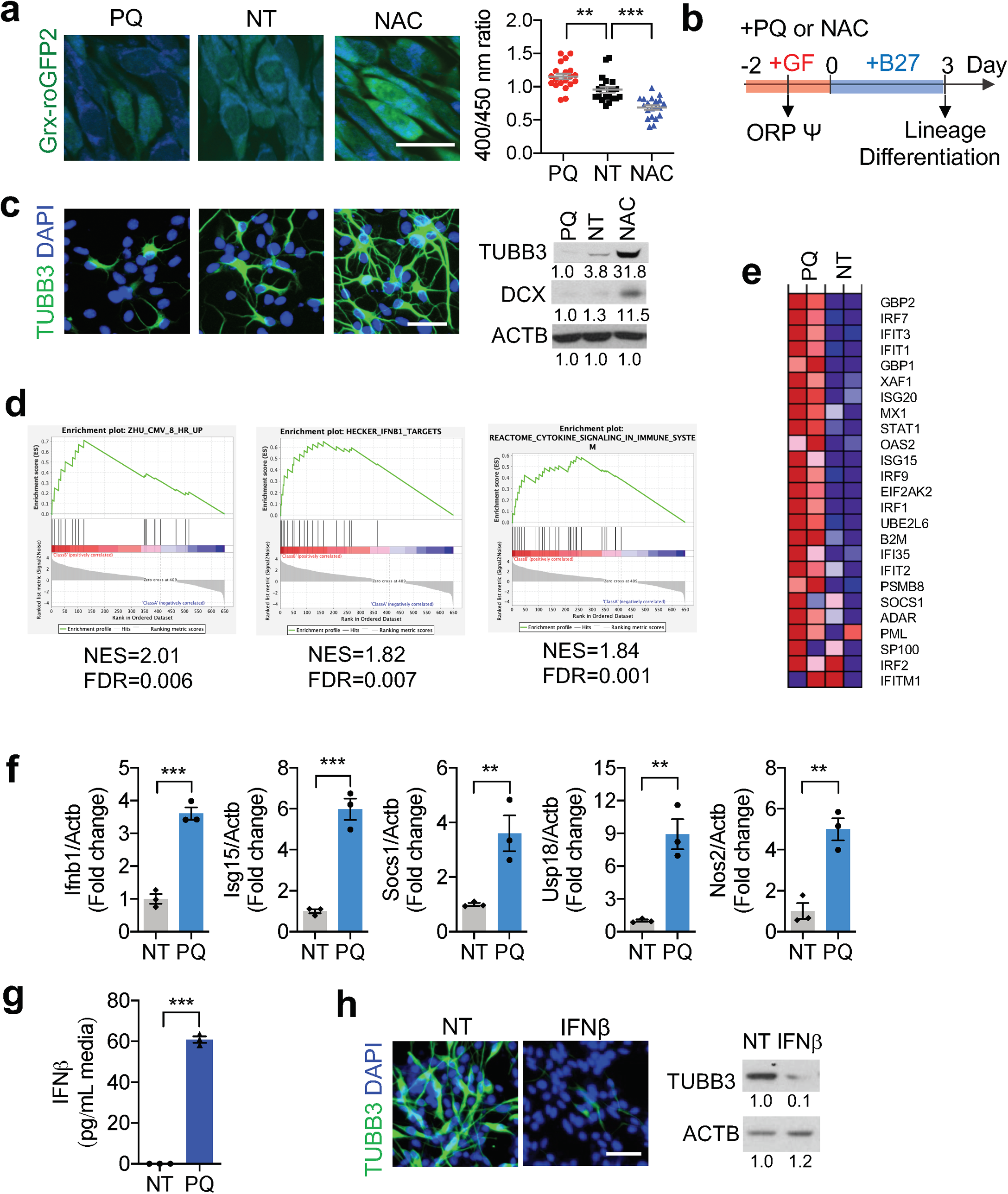
Cellular stress response under high redox potential activates IFN-I responses. **a**. Left, representative images of treated Grx1-roGFP2-NSPC. Right, quantitation for redox potential of Grx1-roGFP2-NSPC following 24 h treatment. Scale bar = 20 µm. Mean ± s.e.m. of 20 cells. PQ: 5 µM of paraquat, NT: mock treated control, NAC: 5 mM of N-acetylcysteine. **b**. Schema for treatment of NSPC culture. ORP: oxidation/reduction potential. **c**. IF (left) and WB (right) analysis for TUBB3 or DCX following 3 days of differentiation. Scale bar = 50 µm. **d**. GSEA of differentially expressed genes following 2 days of PQ treatment in NSPCs. **e**. Heatmap of the gene set from Hecker_IFNB1_Targets. **f.** qRT-PCR results for ISGs after 4 days of PQ treatment. Mean ± s.e.m. of 3 independent experiments. **g.** IFNβ secretion in the media following 48 h treatment. Mean ± s.e.m. of 3 independent experiments. Statistical significance was determined by one-way ANOVA for **a** and by unpaired t-test for **f** and **g**. **P < 0.01; ***P < 0.001. **h.** IF (left) and WB (right) analysis of NSPC differentiated with or without IFNβ (40 ng/ml) treatment for 2 days. Scale bar=50 μm. Experiments for **c** and **h** were repeated three times independently with similar results and representative images/blots are shown.

To determine the signaling pathway that underlies the oxidative stress-induced neurogenic decline, we next performed gene expression profiling against mock-treated control and NSPCs following 48 h redox preconditioning. Gene set enrichment analysis (GSEA) of differentially regulated genes revealed type-I Interferon (IFN-I) signaling as one of the most enriched signature pathways (Fig. 1d, e). Quantitative real-time PCR (qRT-PCR) further confirmed transcriptional upregulation of major IFN-I signaling downstream surrogates, including *Ifnb1, Isg15, Socs1, Usp18, and Nos2* (Fig. 1f). Moreover, ELISA analysis further revealed that secretion of the key IFN-I response effector IFNβ was also strongly elevated in PQ-treated NSPCs (Fig. 1g). To test whether activation of IFN-I signaling accounts for redox stress-induced neurogenic decline, we treated the NSPCs with IFNβ. Indeed, addition of IFNβ alone was sufficient to suppress neurogenic differentiation of NSPCs (Fig. 1h). These data collectively suggest that oxidative stress signaling regulates neurogenic differentiation through IFN-I pathway.

### FOXO3 is required for ROS-induced IFN-I response

FOXO proteins are major regulators of physiological oxidative stress response partly, because they modulate the transcriptional expression of ROS-scavenging enzymes^26, 27^. To determine the role of FOXO3 in ROS-induced IFN-I response, we next analyzed how FOXO3 depletion impacts oxidative stress-induced IFN-I signaling activation. Unsurprisingly, PQ treatment to the control NSPCs (non-targeted guide RNA, sg-NT) elicited a robust IFN-I response, as evidenced by the markedly enhanced phosphorylation of IFN-I upstream and downstream signaling protein Stat1 and TBK1 (Fig. 2a, b), elevated IFNβ secretion (Fig. 2c), and the strongly upregulated mRNA expression of IFN-I stimulated genes (ISGs) (*Isg15, Socs1, Usp18* and *Nos2*) (Fig. 2d). In comparison, the PQ treatment-induced IFN-I response and its upstream and downstream signaling activation were evidently attenuated in PQ treated NSPCs depleted FOXO3 (sg-Foxo3) (Fig. 2b-d), suggesting that FOXO3 plays a crucial role in regulation of ROS-induced IFN-I response.

**Figure 2.**
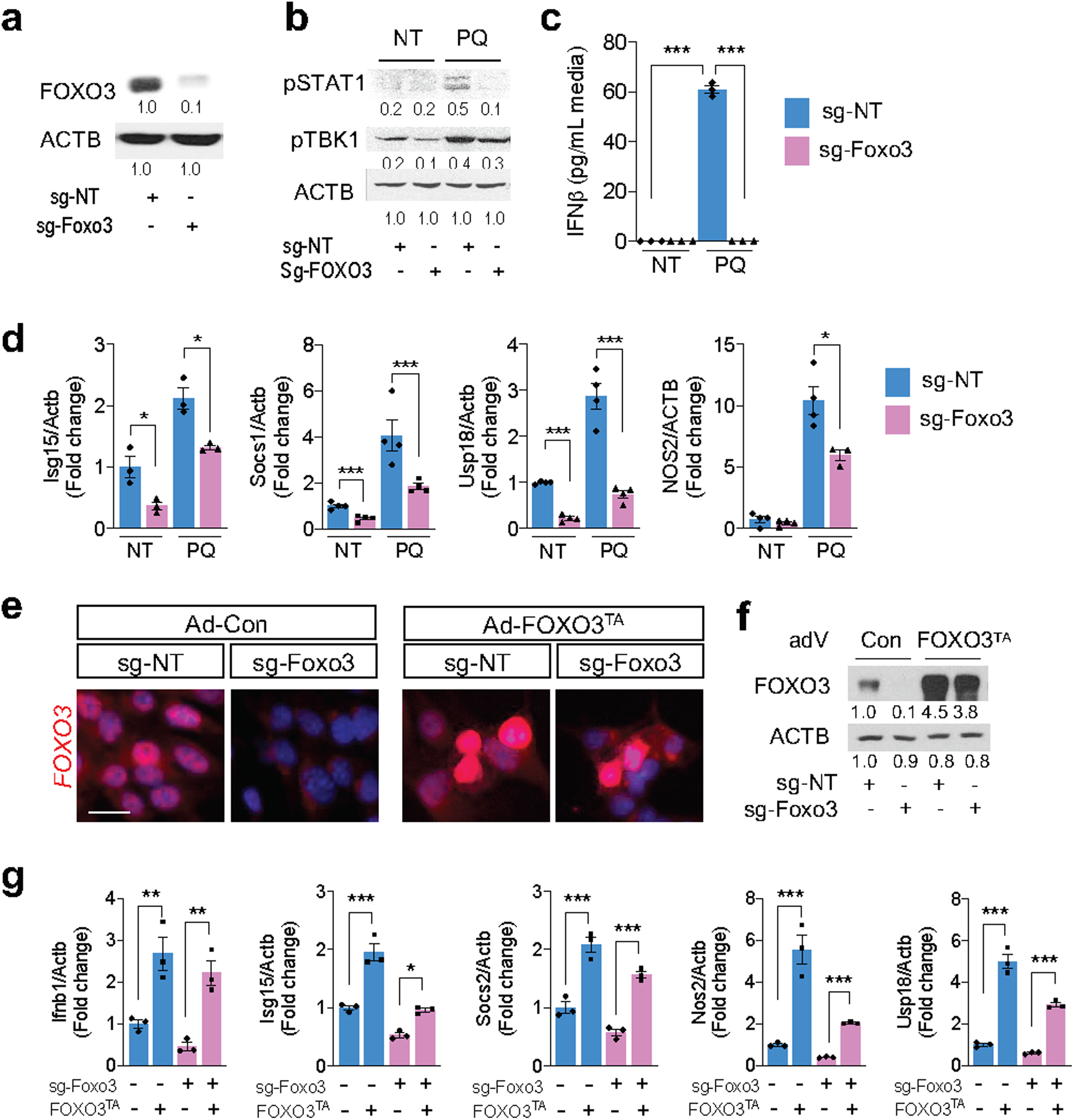
FOXO3 is necessary for ROS-induced IFN-I response. **a**. WB analysis for FOXO3 depletion in sg-Foxo3 infected NSPCs. sg-NT: non-targeted guide RNA infected NSPC, sg-Foxo3: Foxo3-targeted guide RNA infected NSPC. **b**. WB analysis for STAT1 and TBK1 phosphorylation following 48 h treatment. NT: mock treated control, PQ: 5 μM of paraquat. **c**. IFNβ secretion in the media following 48 h treatment. **d**. qRT-PCR results for ISGs following 4 days of PQ treatment. **e** and **f**. IF (**e**) and WB (**f**) analysis for FOXO3 on either adenovirus for control or FOXO3^TA^ infected NSPCs. **g**. qRT-PCR results for ISGs at 4 days after the infection of adenovirus. For **c**, **d**, and **g**, statistical significance was determined by one-way ANOVA. Mean ± s.e.m. of 3 independent experiments. *P < 0.05; **P < 0.01; ***P < 0.001. Experiments for **a, b, e**, and **f** were repeated three times independently with similar results and representative images/blots are shown.

FOXO3 integrates a variety of cellular signals that modulate its transcriptional activity ^28^. To examine whether activation of FOXO3 by itself was sufficient to trigger IFN-I response independently of oxidative stress, we transduced NSPCs with an adenoviral-encoded activated mutant form of FOXO3 (FOXO3^TA^, triple alanine form^29^) that was exclusively located in the nucleus (Fig. 2e, f). qRT-PCR analysis of the FOXO3^TA^ transduced NSPCs revealed a markedly increased expression of ISGs as compared to the control adenovirus-infected NSPCs (Fig. 2g), indicating that FOXO3 is directly responsible for oxidation stress-induced IFN-I activation.

### Oxidation of FOXO3 activates IFN-I response

Previous reports suggest that ROS signaling activates FOXO by inducing its nuclear translocation^30, 31^. Indeed, we found that ROS treatment of NSPCs not only stimulated FOXO3 nuclear retention and activation, but also led to an elevated FOXO3 protein expression (Fig. 3a-c). By contrast, treatment with the antioxidant NAC promoted FOXO3 cytoplasmic shuttling and protein degradation and reduced FOXO3 transcriptional activity. The nucleo-cytoplasmic shuttling of FOXO reportedly is controlled through a combination of post-translational modifications, particularly AKT-mediated phosphorylation that promotes its cytoplasmic sequestering^28^. Consistently, ROS treatment caused a significant reduction of FOXO3 phosphorylation at threonine 32/serine 256, raising a possibility that oxidative stress may induce FOXO3 nuclear translocation by impeding its phosphorylation^32, 33^.

**Figure 3.**
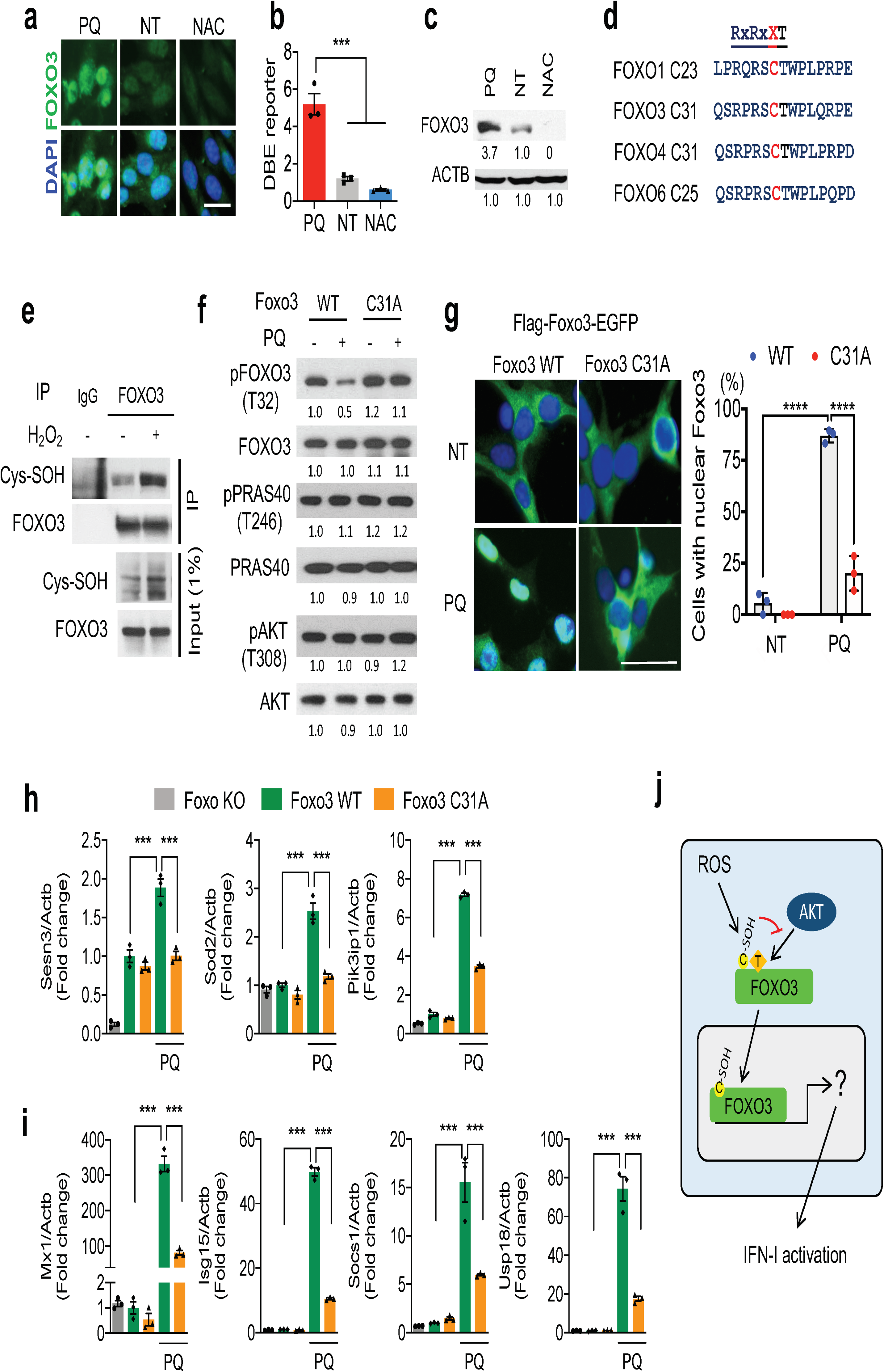
Oxidation at Cys31 of FOXO3 activates IFN-I response. **a-c**. IF (**a**), DBE reporter (**b**), and WB (**c**) analysis for FOXO3 in NSPCs following 24 h of respective treatment. Scale bar = 20 μm. **d**. Conserved consensus sequence adjacent to AKT phosphorylation site of mouse FOXO3 protein. **e**. WB for cysteine sulfenylation (Cys-SOH) following immunoprecipitation of FOXO3. **f**. WB analysis for indicated proteins from non-treated and PQ treated (400 μM, 0.5 h) Foxo3 WT or C31A mutant transduced NSPCs. **g**. Microscopic analysis of Foxo3 WT or C31A mutant with c-terminus EGFP tag with or without PQ treatment (400 μM, 16 h). % of cells with nuclear Foxo3 is plotted on the right. **h** and **i**. qRT-PCR analysis for transcriptional targets of FOXO3 (**h**) and ISGs (**i**). Foxo null NSPCs with WT or C31A mutant Foxo3 were analyzed following 4 days of PQ treatment. j. Schema for activation of FOXO3 by oxidation at Cys31 residue. For **b**, **g**, **h**, and **i**, statistical significance was determined by one-way ANOVA. Mean ± s.e.m. of 3 independent experiments. ***P < 0.001; ****p<0.0001. Experiments for **a, c, e**, and **f** were repeated three times independently with similar results and representative images/blots are shown.

Reversible cysteine thiol oxidation is a well-known mechanism that regulates signaling cascades and protein activities under redox stress conditions^34^. Immunoprecipitation followed by blotting analysis against cysteine sulfenic acid (Cys-SOH) confirmed a strong elevation of FOXO3 sulfenylation in ROS treated NSPCs compared to the controls (Fig. 3e). Since mammalian FOXO3 contains a highly conserved Cys residue (Cys31) adjacent to threonine (Thr) that is subjected to AKT phosphorylation (Fig. 3d), we next asked whether the Cys31 oxidation affected AKT-dependent Thr32 phosphorylation. To this end, we reconstituted the FOXO3-null NSPCs with a lentiviral construct encoding green fluorescent protein (GFP)-tagged either wild-type (WT) or Cys31 to alanine FOXO3 mutant (C31A). Immunoblot analysis showed that ROS treatment induced a strong reduction of Thr32 phosphorylation in FOXO3 WT but not the C31A FOXO3 mutant compared to the mock-treated control cells (Fig. 3f), suggesting that oxidation at Cys31 may impedes Thr32 phosphorylation. In line with this, fluorescence live-imaging of the FOXO3 localization indicated that the ROS-induced nuclear translocation of GFP-tagged FOXO3 C31A mutant was significantly compromised as compared to that of the wild-type FOXO3 (Fig. 3g). Concurrently, qRTPCR analysis of ROS-treated NSPCs revealed upregulation of oxidative stress-induced FOXO3 downstream anti-oxidant genes (i.e. *Sod2, Sesn3*) as well as markedly subdued ISGs in C31A mutant transduced NSPCs as compared to wild-type FOXO3-transduced control cells (Fig. 3h, i). Notably, although C31A mutation compromised ROS-induced FOXO3 nuclear shuttling and activity, it did not affect FOXO3 nuclear translocation upon treatment with either PI3K inhibitor (GDC0941) or AKT inhibitor (MK2206) (Extended Data Fig. 1a, b). These findings together indicate that Cys thiol oxidation and its associated inhibitory function on FOXO3 phosphorylation is a key mechanism that underlies ROS-induced FOXO3 and its downstream signaling activation (Fig. 3j).

### Compromised lamin processing upon oxidative stress invokes IFN-I response

The IFN-I signaling is a cellular innate immune response and is often triggered by the cytosolic DNA-sensing cGAS/STING pathway^35^. To examine whether ROS-induced IFN-I activation is mediated by aberrant cytosolic DNA appearance, we treated the cGAS-GFP-expressing NSPCs with pro-oxidant PQ. Fluorescence microscopic analysis of the PQ treated cells revealed an increased nuclear leakage as represented by appearance of lobulated nuclei and inappropriate formation of cytoplasmic cGAS-GFP-containing DNA foci (Fig. 4a), suggesting a compromised nuclear envelope integrity.

**Figure 4.**
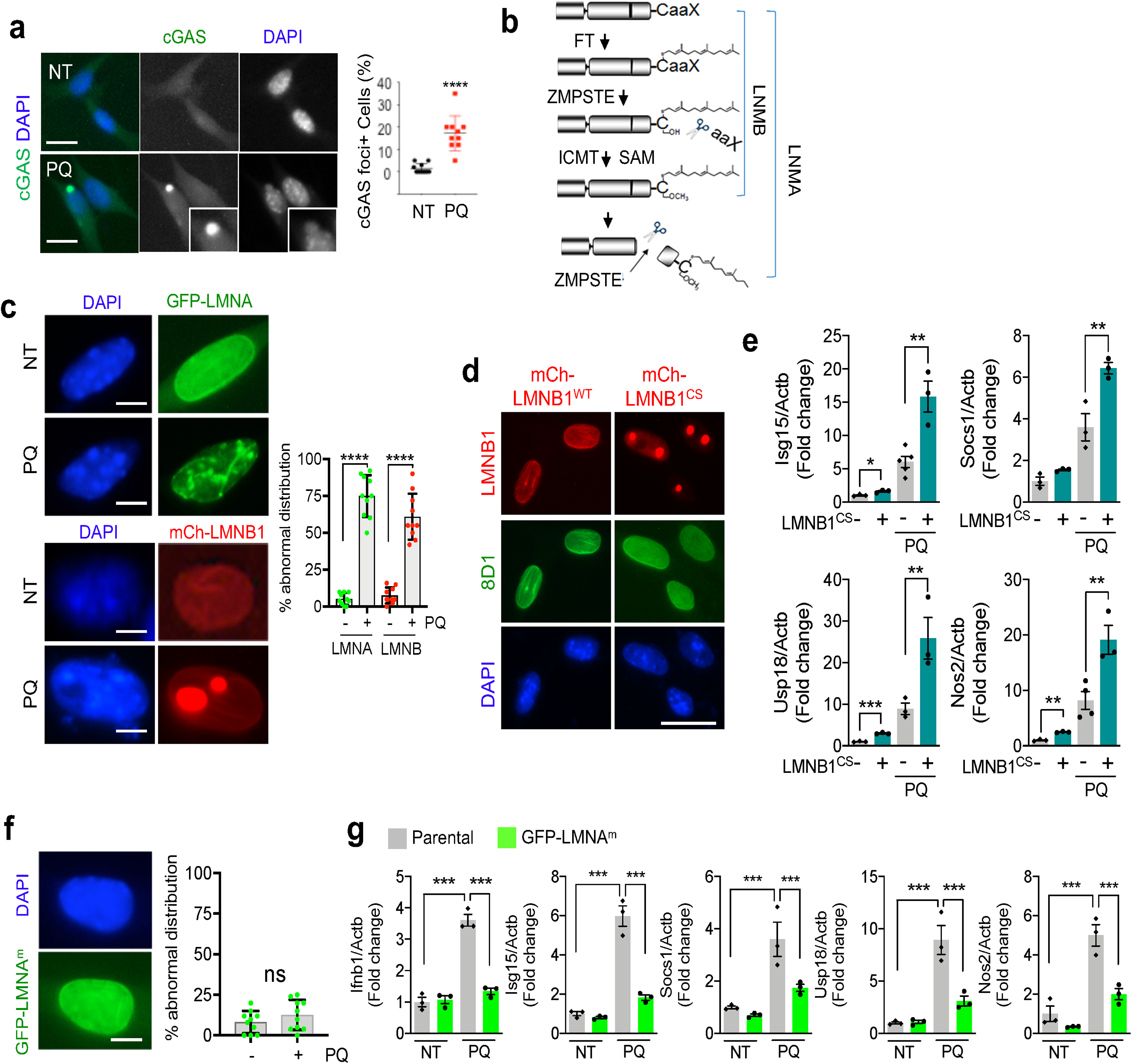
Defective nuclear lamin processing causes nuclear leakage under oxidative stress. **a**. Analysis for cGAS-GFP reporter following 24 h of PQ treatment. Right, quantitation of the percent of cells with cGAS-GFP foci. Scale bar= 4 µm. Mean ± s.e.m. of 10 images. **b**. Diagram for lamin processing. FT: Farnesyl transferase, ZMPSTE: ZMPSTE24 protease, ICMT: Isoprenylcysteine carboxymethyltransferase, SAM: S-adenosyl-L-methionine. **c**. Left, microscopic analysis for GFP-LMNA and mCherry-LMNB1 localization after 24 h of PQ treatment. Right, quantitation for abnormal lamin structure ratio to whole cells. Scale bar= 2 µm. Mean ± s.e.m. of 10 images. **d**. IF analysis for mature LMNB1 (8D1, green) in NSPC expressing mCherry tagged prelamin B1 (mCh-LMNB1^WT^) and mCherry tagged mutant prelamin B1 (mCh-LMNB1^cs^). Scale bar= 5 μm. Experiment was repeated three times independently with similar results and representative images are shown. **e**. qRT-PCR analysis for ISGs after 4 days of PQ treatment. Mean ± s.e.m. of 3 independent experiments. **f.** NSPC expressing mature LMNA (GFP-LMNA^m^). Mean ± s.e.m. of 10 images. **g**. qPCR results for ISGs after 4 day of PQ treatment. Mean ± s.e.m. of 3 independent experiments. Statistical significance was determined by unpaired t-test for **a**, **c**, and **f** and by one-way ANOVA for **e** and **g**. *P < 0.05; **P < 0.01; ***P < 0.001; ****p<0.0001; ns, not significant.

The nuclear lamina is essential for the maintenance of nuclear shape and mechanics, and its dysregulation causes nuclear envelopathies and accumulation of cytosolic chromatin fragments^36^. As an essential component of nuclear lamina, the maturation of functional lamin A/B from newly synthesized prelamin A/B follows a multistep process of posttranslational modification that involves farnesylation and methylation of its C-terminal cysteine before proteolytic cleavage of its C-terminal 15 amino acids (Fig. 4b). To test whether oxidative stress impacts lamin distribution, we stably transduced NSPCs with N-terminal GFP-tagged prelamin A or mCherry-tagged prelamin B1. Strikingly, compared to the control cells in which the tagged lamin proteins dispersed evenly along the nuclear envelopes, we found that a large portion of PQ-treated NSPCs exhibited an irregular lamin distribution, reminiscent of protein aggregation (Fig. 4c). To test whether disrupted lamin processing activated IFN-I response upon oxidative stress, we stably expressed the cysteine 585 to serine prelamin B1 mutant (LMNB1^CS^) in NSPCs, that is defective of prelamin maturation-essential farnesylation and methylation^37^. Immunofluorescence analysis of the cells indicated that LMNB1^CS^ mutant protein, which was negative to mature lamin B1-specific 8D1 monoclonal antibody^38^, formed the aggregate-like nucleoplasmic foci similar to the ones observed in wild-type lamin B1-transduced NSPCs under ROS treatment (Fig. 4d). qRT-PCR analysis further revealed that expression of LMNB1^CS^ mutant alone activated IFN-I response and downstream gene expression and that ROS treatment could further enhance its effect on ISGs expression (Fig. 4e). Conversely, expression of a C-terminus deletion form of mature lamin A mutant (LMNA^m^) strongly attenuated the ROS-induced IFN-I signaling and ISGs expression (Fig. 4f, g). These findings together strongly suggest defective lamin processing as an underlying cause of IFN-I response under oxidative stress.

### ROS-induced intracellular SAM depletion disrupts lamin maturation

Lamin maturation requires isoprenylation and methylation on the c-terminal cysteine residues^39^. To determine how ROS regulates lamin posttranslational modification, we performed targeted quantitative polar metabolomics profiling by liquid chromatography-tandem mass spectrometry (LC-MS) on samples derived from control, pro- or anti-oxidant treated NSPCs. Among the 258 metabolites analyzed, we found that the turnover of SAM exhibited an inverse correlation with redox potential. Compared to the mock-treated control cells, treatment with the pro-oxidant PQ gave rise to a 3.3-fold reduction of cellular SAM and a 1.7-fold reduction of SAM to SAH ratio (Fig. 5a). By contrast, NSPCs treated with anti-oxidant NAC exhibited a 2.5-fold increase of cellular SAM level.

**Figure 5.**
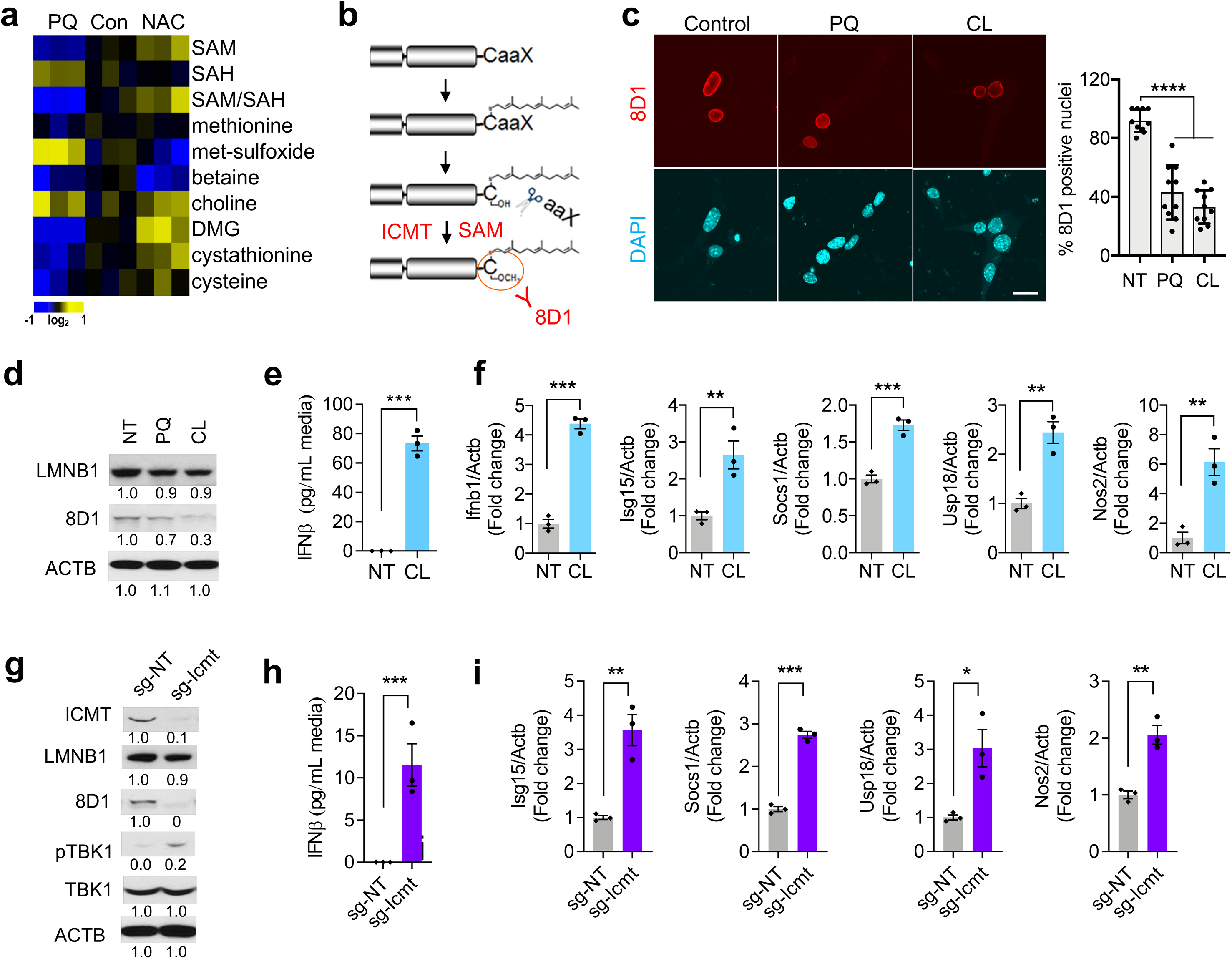
Stress-depleted intracellular SAMe underlies nuclear leakage through lamin B1 maturation failure. **a**. Heat map for metabolites extracted from NSPCs after 2 days of respective treatment (n=3). SAM: S-adenosyl-methionine, SAH: S-adenosyl-homocysteine, DMG: dimethylglycine. **b**. Diagram for lamin processing. **c**. Left, IF analysis for mature laminB1 (8D1, red) following 48 h of PQ or CL treatment. Right, quantitation for 8D1 positive nuclei. Statistical significance was determined by one-way ANOVA. Mean ± s.e.m. of 10 images. ****P<0.0001. **d**. WB analysis for total laminB1 (LMNB1) and mature laminB1 (8D1). **e** and **f**. IFNβ secretion in the media (**e**) and qRT-PCR analysis for ISGs (**f**) after SAM depletion following 48h of CL treatment. CL:1-aminocyclopentanecarboxylic acid**. g**. WB analysis for defect of lamin processing and IFN-I activation in ICMT-targeted guide RNA transduced NSPCs (sg-Icmt). **h** and **i**. IFNβ secretion in the media (**h**) and qRT-PCR analysis for ISGs (**i**) following 48 h growth. For **e**, **f**, **h**, and **i**, statistical significance was determined by unpaired t-test. Mean ± s.e.m. of 3 independent experiments. *P < 0.05; **P < 0.01; ***P < 0.001. Experiments for **d** and **g** were repeated three times independently with similar results and representative blots are shown.

SAM is a principal methyl donor for a variety of biological processes including isoprenylcysteine carboxymethyl transferase (ICMT)-mediated lamin methylation (Fig. 5b). Since prelamin methylation is a prerequisite step of lamin maturation, we next examined the effect of SAM depletion on lamin processing by treating NSPCs with cycloleucine (CL), a methionine adenosyltransferase 2A (MAT2A) inhibitor. Immunofluorescence and Immunoblot analyses revealed that inhibition of SAM production compromised lamin maturation as evidenced by the significantly reduced levels of 8D1-positive mature lamin B1 in CL- or PQ-treated NSPCs compared to the mock controls (Fig. 5c, d). In addition, CL treatment alone was sufficient to elicit IFNβ secretion and induction of ISGs (Fig. 5e, f). Consistently, disrupting lamin methylation by depletion of its methyl transferase ICMT (sg-Icmt) inhibited lamin B1 maturation and provoked IFN-I response, including induction of TBK1 phosphorylation (Fig. 5g and extended data fig. 2a), IFNβ secretion (Fig. 5h), and ISGs expression (Fig. 5i). Together, these findings suggest that SAM depletion and its dependent lamin methylation disruption are the underlying cause of ROS-induced cGAS/STING-IFN-I signaling activation.

### ROS regulates intracellular SAM through GNMT

SAM is a universal co-substrate involved in methyl group transfers^40^. Intracellular SAM levels are balanced by MAT2A-catalyzed synthesis and its consumption through multiple catabolic processes (Fig. 6a). Since our metabolite profiling revealed little change of intracellular methionine - the precursor for SAM (Fig. 6a), we next turned to the expression of the major enzymes involved in SAM metabolism. qRT-PCR analysis of control and ROS-treated NSPCs indicated that cellular expression of MAT2A, the enzyme that catalyzes the synthesis of SAM from methionine, remained relatively stable (Extended Data Fig. 3a). By contrast, among the key catabolic enzymes that catalyze the SAM to SAH conversion, we found that expression of glycine N-methyl transferase (GNMT) was markedly induced by PQ treatment but suppressed by anti-oxidant NAC (Fig. 6b, c).

**Figure 6.**
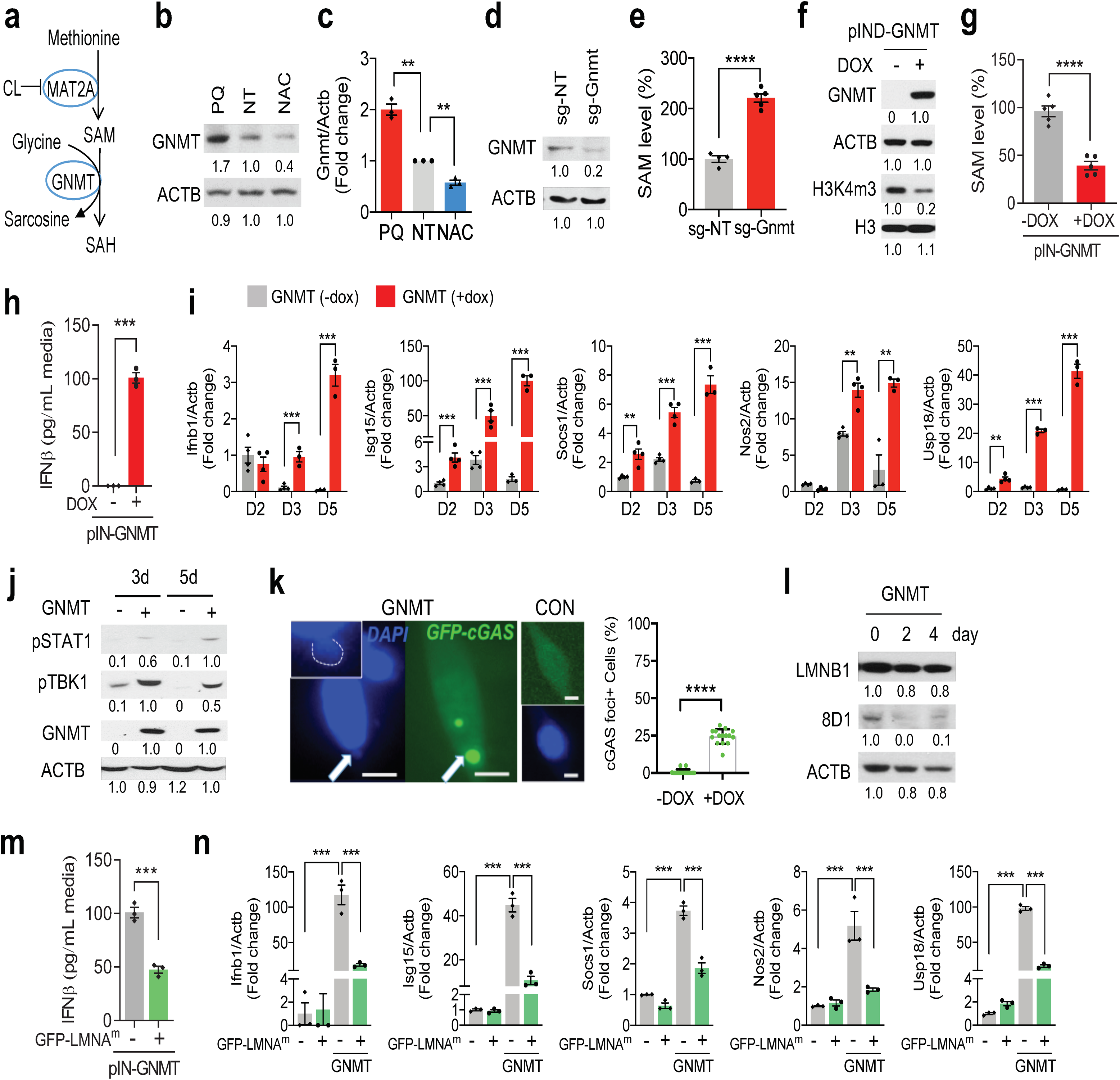
Stress-induced GNMT depletes intracellular SAM. **a**. Diagram for GNMT function. MAT: Methionine adenosyltransferase, GNMT: Glycine N-methyltransferase. **b** and **c**. WB (**b**) or qRT-PCR (**c**) analysis for GNMT expression in NSPCs following 2 days of indicated treatment. Mean ± s.e.m. of 3 independent experiments. **d**. WB analysis for GNMT in GNMT-targeted guide RNA-expressing NSPC (sg-Gnmt). **e**. SAM levels following 48 h treatment in sg-NT (black bar) or sg-Gnmt (red bar) infected NSPC. 100% of SAM = 50 μM. Mean ± s.e.m. of 5 independent experiments. **f** - **h**. WB analysis (**f**), SAM level (n=5) (**g**), and IFNβ secretion in the media (n=3) (**h**) following induction of GNMT with 2 μg/mL of doxycycline (DOX) for 2 days. 100% of SAM = 50 μM. Mean ± s.e.m. of 3-5 independent experiments. **i**. qRTPCR results for ISGs at indicated points after induction of GNMT. Mean ± s.e.m. of 4 independent experiments. **j**. WB analysis for STAT1 or TBK phosphorylation following induction of GNMT at indicated points. **k**. cGAS-GFP reporter analysis after induction of GNMT. Represented number is the result of counting cells with cGAS-GFP foci. Scale bar= 2 μm. Mean ± s.e.m. of 15 images. **l**. WB analysis for total lamin B1(LMNB1) and mature lamin B1 (8D1) expression after induction of GNMT. **m**. IFNβ secretion in the media following 48 h treatment of DOX in GFP-LMNA^m^ expressing NSPCs (GFPLMNA^m^). Mean ± s.e.m. of 3 independent experiments. **n**. qRT-PCR for ISGs following 4 days of GNMT induction. Mean ± s.e.m. of 3 independent experiments. Statistical significance was determined by unpaired t-test for **e, g, h, i, k**, and **m**, and by one-way ANOVA for **c** and **n**. **P0.01; ***P < 0.001; ****P<0.0001. Experiments for b, d, f, j, and i were repeated three times independently with similar results and representative blots are shown.

GNMT catalyzes the reaction of glycine to sarcosine by using SAM as the methyl donor^41^. CRISPR/Cas9-mediated GNMT depletion by guide RNA (sg-Gnmt) led to 2-fold enhancement of cellular SAM accumulation compared to the control sgRNA transduced cells (Fig. 6d, e). Conversely, doxycycline (DOX)-induced overexpression of exogenous GNMT conferred a rapid ~ 70% SAM depletion within 24 h that led to a reduction of global H3K4 methylation (Fig. 6f, g), consistent with its role as a principle methyl donor for histone methylation^42, 43^. Concurrently, DOX-induced GNMT expression was also sufficient to increase IFNβ secretion that initiate a time-dependent IFN-I response and activate its downstream signaling and gene expression (Fig. 6h-j). These findings suggest GNMT-regulated SAM depletion as a likely route to ROS-induced IFN-I activation.

To determine whether GNMT-regulated SAM depletion could also instigate nuclear leakage accompanied by cGAS/STING signaling activation, we transduced the DOX-inducible GNMT expressing NSPCs with the cytosolic DNA fragment-sensing cGAS-GFP construct. Compared to the mock-treated control cells, DOX-treated NSPCs exhibited a significant elevation of cGAS-GFP-containing foci formation (24.4% ± 1.275% vs 0.7% ± 0.45%) (Fig. 6k). Immunoblot analysis of control and DOX-treated NSPCs further revealed a strong reduction of 8D1-positive mature lamin B1 protein level following GNMT induction (Fig. 6l), suggesting compromised lamin maturation. Consistently, transduction of a mature lamin mutant (GFP-LMNA^m^) in the DOX-GNMT NSPCs restored the nuclear envelop integrity and suppressed the GNMT induction-evoked IFNβ secretion as well as ISGs expression (Fig. 6m, n). Concordantly, enforced expression of a STING^HAQ^ mutant44 could also partially offset the effect of GNMT induction by attenuating the IFN-I response and downstream gene expression (Extended Data Fig. 3b, c). Collectively, these data indicate that GNMT is a key regulator of IFN-I response under ROS treatment.

### Redox stress impacts NSPC neurogenic potential through FOXO3-regulated GNMT expression

We observed that the frequency of PQ-induced cGAS-GFP foci was suppressed by FOXO3 depletion (Extended data fig. 4a, b). Consistently, expression of FOXO3^TA^ increased irregular nuclei with decreased 8D1 staining (Extended data fig. 4c, d), phenocopying GNMT induced NSPCs. Given our findings that FOXO3 and GNMT were both involved in ROS-induced IFN-I activation, we next investigated the potential connection between FOXO3 activation and GNMT induction under stress conditions. By surveying the +/- 3 kb genomic DNA sequence near murine GNMT transcription start site (TSS), we identified two putative DAF‐16 family binding element (DBE) FOXO motifs positioned 200-300 bp upstream of TSS (Extended Data Fig. 5). Chromatin immunoprecipitation (ChIP) coupled with qRT-PCR confirmed that FOXO3 was 9.7 ± 2.7-fold enriched at GNMT promoter relative to the background gene desert (Fig. 7a). Moreover, qRT-PCR analysis of NSPCs indicated that relative to control virus infected NSPCs, enforced expression of an active FOXO3^TA^ mutant was able to significantly enhance GNMT transcription (Fig. 7b), whereas CRISPR/Cas9-mediated depletion of endogenous FOXO3 suppressed GNMT mRNA expression (Fig. 7c). These data suggest that FOXO3 transcriptionally controls GNMT expression.

**Figure 7.**
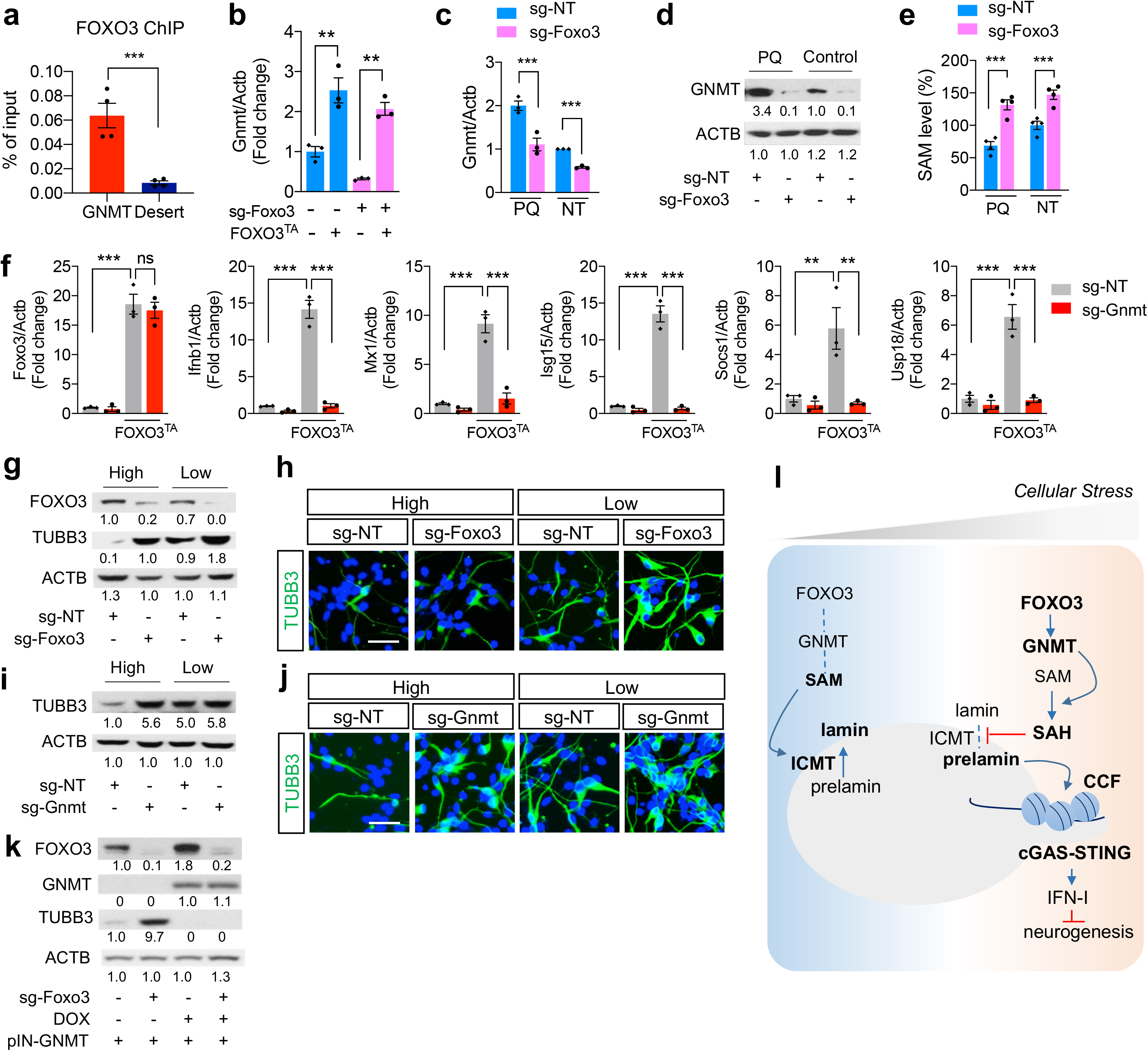
Redox stress impacts neurogenic potential of NSPC through regulating SAMe levels. **a**. FOXO3 ChIP-qPCR analysis at *Gnmt* promoter in comparison to a gene desert region. Mean ± s.e.m. of 4 independent experiments. **b** - **d**. qRT-PCR analysis for Gnmt mRNA expression (**b** and **c**) and WB analysis for GNMT expression (**d**) of NSPCs treated with PQ for 48 h. Mean ± s.e.m. of 3 independent experiments. **e**. SAM levels in sg-NT vs. sg-Foxo3 NSPCs treated for 48 h as indicated. 100% of SAM = 50 μM. Mean ± s.e.m. of 4 independent experiments. **f.** qRT-PCR results for Foxo3 and ISGs on either control adenovirus or FOXO3-TA adenovirus infected NSPC. Mean ± s.e.m. of 3 independent experiments. Statistical significance was determined by unpaired t-test for **a** and by one-way ANOVA for **b, c, e**, and **f**. *P < 0.05; **P < 0.01; ***P < 0.001. **g-j**. WB (**g**, **i**, and **k**) and IF (**h** and **j**) analysis for TUBB3 expressions each NSPC line at 3 days of differentiations. Scale bar= 40 μm. Experiments for d, g, h, i, j, and k were repeated three times independently with similar results and representative images/blots are shown. **k**. Model for the mechanism how cellular stress elicits IFN-I response and inhibits neurogenic differentiation potential of neural stem cells.

We next examined whether FOXO3 regulated ROS-initiated IFN-I response through GNMT. Treatment of NSPCs with pro-oxidant agent PQ promoted a marked increase of GNMT mRNA and protein expression relative to the mock-treated control cells (Fig. 7c, d). This ROS-induced GNMT upregulation was significantly compromised in the NSPCs depleted of FOXO3. Notably, the FOXO3-depleted NSPCs exhibited a steady increase of cellular SAM levels compared to their respective controls before or after ROS treatment (Fig. 7e). Concordantly, knockdown of GNMT abolished FOXO3^TA^ expression-induced IFN-I response and downstream gene expression (Fig. 7f).

Finally, we went on to determine how FOXO3- GNMT/SAM-IFN-I signaling pathway regulates neurogenesis under oxidative stress condition. As expected, immunoblot and immunofluorescence analysis revealed that treatment of NSPCs with pro-oxidant PQ attenuated their neuronal differentiation capacity, as evidenced by the reduction of expression of neuronal marker TUBB3 and percentage of TUBB3-positive cell population relative to the mock-treated control cells (Fig. 7g-j). But further depletion of either FOXO3 or GNMT in the PQ-treated NSPCs reversed the ROS effect and was sufficient to restore their neurogenic potential (Fig. 7g-j). Notably, the FOXO3 depletion-promoted neuronal differentiation could be further blocked by DOX-induced exogenous GNMT expression, consistent with our finding that GNMT is a downstream effector of FOXO3 signaling (Fig. 7k). Considering the elevation of ROS in aging brain, we further examined type-I IFN stimulated gene expression in young and old (<60 year-old) and aged (>60 year-old) patient brain samples. Consistently, we observed a clear increase of ISGs along with GNMT mRNA expression in aged brains (Extended Fig. 6). Altogether, our results suggest the FOXO3-GNMT/SAM-lamin-cGAS/STING-IFN-I signaling cascade as an important physiological stress response program that may protect the nervous system against acute oxidative insults (Fig. 7l).

## Discussion

Alterations of the redox state, as in many brain pathologies, regulate the fate of NSPCs^45^. Our study revealed that cellular stresses including a higher redox potential are translated into IFN-I response via FOXO3-GNMT/SAM-lamin changes (Fig. 7l). In particular, we showed that redox potential controls NSPC function by altering IFN-I response through metabolic regulation of intracellular SAM availability. Mechanistically, our study uncovered a previously unidentified FOXO3 signaling cascade that functionally connects oxidative stress response with NSPC differentiation through SAM-depletion-induced IFN-I activation. Our findings of redox-dependent neurogenic regulation warrant future studies on the therapeutic rejuvenation of stress-impacted adult NSPCs.

FOXO transcription factors play a central role in a wide range of biological processes, including stress sensing and regulation of stress response^15^. Genetic studies from many organisms have repeatedly demonstrated the conserved insulin/IGF-PI3K-AKT-FOXO cascade as a major regulatory signaling pathway of aging and lifespan. In the central nervous system, expression of FOXO plays not only a key role in preserving neural stem cell pools^9, 10^, but also protects neurons against age-related axonal degeneration across species^16–18^. Despite these advances, there still lacks a mechanistic understanding of how oxidative stress affects FOXO activation systematically and whether and how that contributes to the neuroprotective responses. In the current study, we identified FOXO3 oxidation at the evolutionarily conserved Cys31 residue as a new regulatory mechanism that modulate redox-dependent FOXO3 nucleo-cytoplasmic shuttling and downstream signaling. Notably, a previous study reported that ROS-induced FOXO4 oxidation at Cys239 promotes its nuclear import by forming a disulfide-dependent protein complex with transportin-1^31^. These findings suggest that redox-regulated nuclear shuttling is a conserved mechanism underlying FOXO-mediated oxidative stress response.

Our data indicate that FOXO3 mediates redox response through regulation of GNMT and downstream SAM catabolism. Enhanced SAM catabolism by GNMT extends the lifespan in *Drosophila*^46^. In the nervous system, GNMT-mediated SAM metabolism is required for the proliferative signaling of NSPC and hippocampal neurogenesis^47^. But the underlying mechanism is unclear. Here we found that treatment of NSPCs with prooxidants led to upregulated GNMT expression and reduction of intracellular SAM availability. SAM is a metabolite generated via the one-carbon metabolism and is the main methyl donor in cellular methylation reactions^40^. SAM depletion through dietary methionine restriction has been shown to modulate histone methylation and induce stem cell quiescence^42, 43, 48^. In our study, we found that not only GNMT-induced SAM depletion in NSPCs confers a global reduction of H3K4 methylation, but is also sufficient to trigger cGAS/STING signaling and IFN-I response through regulation of nuclear lamin maturation. These findings support FOXO3-GNMT/SAM axis as a stress responsive program that protect tissue homeostasis by orchestrating anti-oxidative function, metabolic rewiring, and gene expression.

Defective lamin processing is known to cause various human pathologies, particularly those related to aging. A truncated lamin A causes a premature aging syndrome of Hutchinson-Gilford progeria. Consistent with our findings, expression of mutant lamin activated IFN-I response^49^. Additionally, recent reports suggest that lamins play important roles in both the outside-in and inside-out signaling processes. External mechanical forces trigger changes in nuclear envelope structure and composition, chromatin organization and gene expression^50^. Likewise, lamin A is stabilized upon external stress to protect the genome^51^. These studies agree with our findings linking lamin and cellular stress response. We found that FOXO3 mediates oxidative stress response through regulation of intracellular SAM availability and nuclear lamin posttranslational modification. Introduction of a C-terminus deletion form of mature lamin A mutant (LMNA^m^) blunts ROS-induced activation of cGAS/STING signaling and IFN-I response. These findings suggest a new role of nuclear lamin as a signal transducer that mediates oxidative stress response.

Our study contributes to accumulating literature that links cytosolic DNA-sensing cGAS/STING-IFN-I program to aging- and injury-regulated CNS homeostasis. While initially recognized for its critical function in the innate immune response against viral infections, recent studies indicate that cGAS/STING-IFN-I pathway also mediates many other stress responses including signaling from DNA damage and oxidative stresses^52–55^. Notably, increased IFN-I response suppresses proliferation of NSPC and reduces their neuronal differentiation under oxidative stress^13, 24, 56^. In this study, we demonstrated that oxidative stress response activates FOXO3-GNMT/SAM-cGAS/STING-IFN-I signaling cascade and regulates neurogenic potential of NSPCs. Considering increased IFN-I response with declined neurogenesis is an indicative of aging brains^6, 14^, we propose cGAS/STING-IFN-I response as an intrinsic cellular surveillance system that protects NSPCs against the deleterious consequences of oxidative insults.

Consistent with our findings, previous aging studies reported that lowering systemic SAM levels by dietary restriction of its precursor methionine was effective toward extending life span and improving tissue functions in mammals^57^. Engaging FOXO3-GNMT/SAM-lamin-IFN-I response to acute stress conditions is likely to protect organisms against losing long term regenerative potential. This protective mechanism, nevertheless, may drive stem cell dysfunction by increasing quiescence and decreased differentiation potential at the face of chronic pathological stress stimuli. Altogether, our findings revealed novel molecular mechanisms that outline how oxidative stress may trigger IFN-I response-mediated cellular protective response and homeostasis under pathophysiological conditions.

## Methods

### NSPC culture and differentiation

Primary murine NSPC isolated from subventricular zones (SVZ) and cultured as neurospheres are heterogeneous populations with limited repopulation potential. To avoid passage-dependent drift in NSPC populations, we utilized a neonate-derived immortalized *Ink/Arf^−/-^* NSPC culture that maintains the multi-lineage differentiation capability^58^. It contains a mixed population of relatively quiescent neural stem cells (qNSCs), activated NSCs and lineage-committed neuronal precursor cells (NPCs), as well as oligodendroglial progenitors (OPC) based on mRNA expression of lineage markers. NSPC were cultured with N2 media including 20 ng/mL of EGF and bFGF in the presence or absence of 5 mM NAC, 5 μM paraquat, or 40 ng/ml Interferon-β. After 2 days, all growth factors and chemicals were removed and changed to N2 media including B27 supplement to induce differentiation. Cells were harvested at indicated time points for analysis. All the sources of materials are listed on supplementary table.

### Generation of viral particles

Each sgRNA was cloned into lentiCrisprV2 vector following Zhang lab instructions^59^. In brief, annealed sgRNA oligos with T4 PNK enzyme was cloned into lentiCrisprV2 vector digested by BsmBI. pDONR221-GNMT was cloned into pInducer gateway destination vector by using LR clonase II enzyme mix. To generate lentivirus, 1.5 × 10^7^ 293T cells in 150 mm tissue culture dishes were transfected with 18 μg of each plasmid DNA along with 4.5 μg of pMD2.G and 9 μg of psPAX2 packaging vectors using polyethylenimine. The medium containing lentiviral particles were collected at 48 and 72 h after transfection. The expression of GFP-FOXO3^TA^ by adenoviral transduction was performed by incubating NSPC culture with pfu of purified adenoviral particles for 16 hrs. Empty adenoviral particles were used at the same pfu in control cultures. All the sources of plasmid DNAs, materials, viruses, and all oligo sequences are listed on supplementary table.

### Measurement of ROS and redox potential

Intracellular glutathione redox potential was determined by expressing pLPCX cyto Grx1-roGFP2^60^. Grx1-roGFP2 NPCs were treated with 10 mM N-ethylmaleimide for 5 min before fixing with 4% paraformaldehyde to prevent further oxidation. Cells from random fields were scanned by Olympus FLUOVIEW laser scanning confocal microscope using excitation at 405 nm/488 nm. Image analysis was performed using ImageJ software to calculate 405/488 nm ratio.

### Measurement of IFNβ

The medium was collected after the cells were cultured for 2 days. The cells were lysed with laemmli buffer and protein concentrations were determined by BCA assay. IFNβ was measured by using VeriKineTM Mouse IFNβ ELISA kit following the manufacturer’s instructions. All the sources of materials are listed on supplementary table.

### SAM assay

Metabolites were extracted by using cold 80 % methanol from 3 million cells overnight at −80 °C. Relative SAM levels were determined by using MLL1 SAMe-Screener Assay kit following the manufacturer’s instructions. In brief, all standards and samples were incubated with MLL1 enzyme for 15 minutes and subsequently SAM-binding site probe for 15 minutes at room temperature. The levels of free probe for each well were determined by a plate reader (SpectraMax M4) with excitation and emission wavelengths of 575 nm and 620 nm, respectively. All materials are listed on supplementary table.

### RNA extraction, qRT-PCR analysis

Total RNAs were extracted from cells by using NucleoSpin RNA kit. Reverse transcription was carried out on 500 ng of total RNA using utilizing RevertAid RT kit. RT-qPCR was performed on cDNA samples using the PowerUp™ SYBR® Green Master Mix on the 7500 Fast Real-time PCR system. All samples were run in duplicate and the mRNA level of each sample was normalized to that of ACTB mRNA. The relative mRNA level was presented as unit values of 2^dCt (=Ct of ACTB-Ct of gene). All the sources of materials and primers are listed on supplementary table.

### RNA-seq and Data Analysis

Total RNAs were isolated from NPCs treated with or without 5 µM of PQ for 48 hours and subjected to RNA sequencing at the Genomics Resources Core facility of Weill Cornell Medicine. RNA-seq libraries were prepared using the Illumina TruSeq stranded mRNA library preparation kit and sequenced on HiSeq4000 sequencer (Illumina). RNA-seq data were aligned to the mm9 reference genome using STAR 2.3.0e^61^. Raw counts of each transcript were measured by featureCounts v1.4.6-p5^62^. Lists of differentially expressed genes were generated by DESeq2-1.4.5 in R^63^. GSEA analysis in this manuscript were generated from GSEA preranked model. The input of GSEA analysis is the gene expression level logFC (*PQ* versus control).

### Immunofluorescent analysis

Cells were fixed with 4 % paraformaldehyde for 10 min at room temperature followed by permeabilization with 0.2 % triton X-100 in PBS. The cells were subjected to immunofluorescence staining with anti-β-III tubulin (TUBB3, 1:300), anti-lamin B1 (1:1,000), 8D1 (1:500), or FOXO3 (1:1,000) antibodies overnight at 4°C. The cells were then washed with cold PBS, and incubated with Alexa 488-labeled anti-mouse, Alexa 488-labeled anti-rabbit, or Alexa 594-labeled anti-mouse secondary antibodies at room temperature for 1 h. Images were acquired by fluorescence microscopy with EVOS FL Cell Auto Imaging System. All the sources of antibodies and materials are listed on supplementary table.

### Western blot analysis

Cells were lysed by laemmli buffer followed by sonication (30 watt/5 sec/10 cycles). Protein concentration was determined by using Pierce^TM^BCA protein assay kit. 5-30 μg of proteins were fractionated by SDS-PAGE electrophoresis and transferred to PVDF membrane using a transfer apparatus following manufacturer’s instructions. After incubation with 5% skim milk in TBST (10 mM Tris, pH 8.0, 150 mM NaCl, 0.5% Tween 20) for 1 h, the membrane was incubated with antibodies against β-actin (ACTB, 1:10,000), βIII Tubulin (1:1,000), GFAP (1:5,000), H3K4m3 (1:8,000), Histone H3 (1:10,000), GNMT (1:500), FOXO3 (1:1,000), p-FOXO1/3a (T24/T32, 1:1,000), p-TBK1 (S172, 1:1,000), p-STAT1 (1:1,000), p-p65 (1:1,000), p-Akt (S472, 1:2,000), Akt (1:5,000), p-PRAS40 (T246, 1:1,000), ICMT (1:1,000), lamin B1 (8D1, 1:500), or lamin B1 (1:5,000) overnight at 4°C. Membranes were washed three times with TBST for 30 min and then incubated with HRP conjugated anti-mouse or anti-rabbit diluted in 3% skim milk for 1 hour. Blots were washed with TBST three times and developed with the SuperSignal™ West Pico Chemiluminescent substrate according to the manufacturer’s protocols. All the sources of materials and antibodies are listed on supplementary table.

### Immunoprecipitation for detecting Cys-sulfenylation

Cells were pretreated with 2 μM of dinamedone for 1 hr and then, treated with or without 400 μM of PQ for 0.5 hr. Cell were lysed using 1 % TNT buffer (135 mM NaCl, 20 mM Tris-HCl, 1 mM EDTA, and 1% Triton X-100) for 30 min on ice. After centrifugation to remove the debris, 1 mg of protein was incubated with 3 μg of antibody at 4 °C overnight. 15 μl of Dynabeads® β Protein G was added and incubated 4 °C for 3 h. The beads were washed 3 times with 1 % TNT buffer and the proteins were eluted with 2X Laemmli buffer. The protein interaction was determined by western blot.

### Chromatin Immunoprecipitation (ChIP)

ChIP analysis was performed using truChIP Chromatin Shearing Reagent kit. In brief, 3×10^7^ cells were cross-linked for 5 min with 1% paraformaldehyde and quenched with 125 mM glycine for 5 min at room temperature. After nuclei isolation, the chromatin was sheared in shearing buffer (50 mM Tris-HCl, 10 mM EDTA, and 0.1% SDS) using the Covaris M220 focused-ultrasonicator according to the manufacturer’s instructions. Immunoprecipitation was performed with 10 μg of FOXO3 overnight at 4°C. 30 μl of pre-cleared Dynabeads® Protein G was added and incubated for 3 h at 4°C. The beads were washed by high salt buffer (50 mM HEPES, 500 mM NaCl, 1mM EDTA, 0.1% SDS, 1% Triton X-100, and 0.1% sodium deoxycholate) and RIPA buffer (including LiCl) and eluted with elution buffer (50 mM Tris-HCl, 10 mM EDTA, and 1% SDS). After RNase and Proteinase K treatment, eluted DNA was reverse-crosslinked by 65°C incubation overnight. DNA was extracted using NucleoSpin Gel and PCR clean-up DNA extraction kit and size-selection was carried out to obtain <400 bp size DNA fragments using SPRIselect Reagent. qRT-PCR was performed using specific primers. All the sources of materials and antibodies are listed on supplementary table.

### Metabolomics analysis

Ten million cells were homogenized in cold 80 % methanol using homogenizer. Metabolites were extracted over 3 h at −80 °C. Samples were then centrifuged at 4 °C for 15 minutes at 14,000 rpm. The supernatants were extracted and normalized based on tissue weight. Targeted LC/MS analyses were performed on a Q Exactive Orbitrap mass spectrometer coupled to a Vanquish UPLC system. The Q Exactive operated in polarity-switching mode. A Sequant ZIC-HILIC column (2.1 mm i.d. × 150 mm) was used for separation of metabolites. Flow rate was set at 150 μL/min. Buffers consisted of 100% acetonitrile for mobile A, and 0.1% NH4OH/20 mM CH3COONH4 in water for mobile B. Gradient ran from 85% to 30% A in 20 min followed by a wash with 30% A and re-equilibration at 85% A. Metabolites were identified on the basis of exact mass within 5 ppm and standard retention times. Relative metabolite quantitation was performed based on peak area for each metabolite. All data analysis was done using in-house written scripts.

### Statistical analysis

We determined experimental sample sizes on the basis of preliminary data. All results are expressed as mean ± s.e.m. GraphPad Prism software (version 7, San Diego, CA) was used for all statistical analysis. Normal distribution of the sample sets was determined before applying unpaired Student’s two-tailed t-test for two group comparisons. One-way ANOVA was used to assess the differences between multiple groups. The mean values of each group were compared by the Bonferroni’s post-hoc procedure. Differences were considered significant when P<0.05.

### Data availability

The datasets produced in this study are available in the following databases: RNA-seq data: NCBI GEO: GSE146243

## Acknowledgements

Authors thank Dr. Loren Fong (UCLA) for kindly providing prelamin antibodies and discussion. We also thank Drs. Zhe Cheng, Guoan Zhang at the Proteomics and Metabolomics and Tuo Zhang at the Genomics Resources Core Facility of The Weill Cornell Medicine for analysis. This work was supported by the Irma T. Hirschl Award, the Anna Maria and the Kellen foundation, and National Institutes of Health Grant AG048284 (to J. P.).

## Author Information

### Affiliations

*Department of Pathology and Laboratory medicine, Weil Cornell Medicine, New York, NY10021, USA*

Inah Hwang, Teresa Sanchez, Hongwu Zheng, and Jihye Paik

*Department of Medicine, Weil Cornell Medicine, New York, NY10021, USA*

Lewis L Cantley

*Meyer Cancer Center, Weill Cornell Medicine and New York Presbyterian Hospital, New York, NY10021, USA*

Lewis L Cantley and Jihye Paik

*Department of Pharmacology and Cancer Biology, Duke University School of Medicine, Durham, NC 27710, USA*

Ziwei Dai and Jason W Locasale

*Department of Neurosurgery, Southwest Hospital, Chongqing, 400038, China*

Fei Li

## Author Contributions

Conception and design: I.H., H.Z., J.P.

Development of methodology: I.H., H.Z.

Acquisition of data: I.H., J.P.

Analysis and interpretation of data (e.g., statistical analysis, biostatistics, computational analysis): I.H., Z.D., J.W.L., J.P.

Writing, review, and/or revision of the manuscript: I.H., H.Z., J.P.

Administrative, technical, conceptual or material support (i.e., reporting or organizing data, constructing databases): I.H., J.W.L., L.L.C., H.Z., J.P.

Study supervision: J.W.L., T.S., J.P.

## Ethics declarations

LCC is a founder and member of the BOD of Agios Pharmaceuticals and is a founder and receives research support from Petra Pharmaceuticals. These companies are developing novel therapies for cancer. Other authors have declared that no conflict of interest exists.

## References

1. Rossi, D.J., Jamieson, C.H. & Weissman, I.L. Stems cells and the pathways to aging and cancer. Cell 132, 681–696 (2008).

2. Molofsky, A.V. et al. Increasing p16INK4a expression decreases forebrain progenitors and neurogenesis during ageing. Nature 443, 448–452 (2006).

3. Baker, D.J. & Petersen, R.C. Cellular senescence in brain aging and neurodegenerative diseases: evidence and perspectives. J Clin Invest 128, 1208–1216 (2018).

4. Dong, C.M. et al. A stress-induced cellular aging model with postnatal neural stem cells. Cell Death Dis 5, e1116 (2014).

5. Ito, K. et al. Regulation of oxidative stress by ATM is required for self-renewal of haematopoietic stem cells. Nature 431, 997–1002 (2004).

6. Miyamoto, K. et al. Foxo3a is essential for maintenance of the hematopoietic stem cell pool. Cell Stem Cell 1, 101–112 (2007).

7. Tothova, Z. & Gilliland, D.G. FoxO transcription factors and stem cell homeostasis: insights from the hematopoietic system. Cell Stem Cell 1, 140–152 (2007).

8. Yalcin, S. et al. Foxo3 is essential for the regulation of ataxia telangiectasia mutated and oxidative stress-mediated homeostasis of hematopoietic stem cells. J Biol Chem 283, 25692–25705 (2008).

9. Renault, V.M. et al. FoxO3 regulates neural stem cell homeostasis. Cell Stem Cell 5, 527–539 (2009).

10. Paik, J.H. et al. FoxOs cooperatively regulate diverse pathways governing neural stem cell homeostasis. Cell Stem Cell 5, 540–553 (2009).

11. Chuikov, S., Levi, B.P., Smith, M.L. & Morrison, S.J. Prdm 16 promotes stem cell maintenance in multiple tissues, partly by regulating oxidative stress. Nat Cell Biol 12, 999–1006 (2010).

12. Uttara, B., Singh, A.V., Zamboni, P. & Mahajan, R.T. Oxidative stress and neurodegenerative diseases: a review of upstream and downstream antioxidant therapeutic options. Curr Neuropharmacol 7, 65–74 (2009).

13. Kalamakis, G. et al. Quiescence Modulates Stem Cell Maintenance and Regenerative Capacity in the Aging Brain. Cell 176, 1407–1419.e1414 (2019).

14. Zipp, F. & Aktas, O. The brain as a target of inflammation: common pathways link inflammatory and neurodegenerative diseases. Trends Neurosci 29, 518–527 (2006).

15. Martins, R., Lithgow, G.J. & Link, W. Long live FOXO: unraveling the role of FOXO proteins in aging and longevity. Aging Cell 15, 196–207 (2016).

16. Calixto, A., Jara, J.S. & Court, F.A. Diapause formation and downregulation of insulin-like signaling via DAF-16/FOXO delays axonal degeneration and neuronal loss. PLoS Genet 8, e1003141 (2012).

17. Hwang, I. et al. FOXO protects against age-progressive axonal degeneration. Aging Cell 17 (2018).

18. Kaletsky, R. et al. The C. elegans adult neuronal IIS/FOXO transcriptome reveals adult phenotype regulators. Nature 529, 92–96 (2016).

19. Takeuchi, O. & Akira, S. Pattern recognition receptors and inflammation. Cell 140, 805–820 (2010).

20. Ishikawa, H. & Barber, G.N. STING is an endoplasmic reticulum adaptor that facilitates innate immune signalling. Nature 455, 674–678 (2008).

21. Okabe, Y., Sano, T. & Nagata, S. Regulation of the innate immune response by threonine-phosphatase of Eyes absent. Nature 460, 520–524 (2009).

22. Sun, W. et al. ERIS, an endoplasmic reticulum IFN stimulator, activates innate immune signaling through dimerization. Proc Natl Acad Sci U S A 106, 8653–8658 (2009).

23. Schoggins, J.W. et al. A diverse range of gene products are effectors of the type I interferon antiviral response. Nature 472, 481–485 (2011).

24. Baruch, K. et al. Aging. Aging-induced type I interferon response at the choroid plexus negatively affects brain function. Science 346, 89–93 (2014).

25. Monje, M.L., Toda, H. & Palmer, T.D. Inflammatory blockade restores adult hippocampal neurogenesis. Science 302, 1760–1765 (2003).

26. Kops, G.J. et al. Forkhead transcription factor FOXO3a protects quiescent cells from oxidative stress. Nature 419, 316–321 (2002).

27. Nemoto, S. & Finkel, T. Redox regulation of forkhead proteins through a p66shc-dependent signaling pathway. Science 295, 2450–2452 (2002).

28. Calnan, D.R. & Brunet, A. The FoxO code. Oncogene 27, 2276–2288 (2008).

29. Ramaswamy, S., Nakamura, N., Sansal, I., Bergeron, L. & Sellers, W.R. A novel mechanism of gene regulation and tumor suppression by the transcription factor FKHR. Cancer Cell 2, 81–91 (2002).

30. Essers, M.A. et al. FOXO transcription factor activation by oxidative stress mediated by the small GTPase Ral and JNK. EMBO J 23, 4802–4812 (2004).

31. Putker, M. et al. Redox-dependent control of FOXO/DAF-16 by transportin-1. Mol Cell 49, 730–742 (2013).

32. Luo, H. et al. PTEN-regulated AKT/FoxO3a/Bim signaling contributes to reactive oxygen species-mediated apoptosis in selenite-treated colorectal cancer cells. Cell Death Dis 4, e481 (2013).

33. Brunet, A. et al. Stress-dependent regulation of FOXO transcription factors by the SIRT1 deacetylase. Science 303, 2011–2015 (2004).

34. Paulsen, C.E. et al. Peroxide-dependent sulfenylation of the EGFR catalytic site enhances kinase activity. Nat Chem Biol 8, 57–64 (2011).

35. Cai, X., Chiu, Y.H. & Chen, Z.J. The cGAS-cGAMP-STING pathway of cytosolic DNA sensing and signaling. Mol Cell 54, 289–296 (2014).

36. Ivanov, A. et al. Lysosome-mediated processing of chromatin in senescence. J Cell Biol 202, 129–143 (2013).

37. Jung, H.J. et al. Farnesylation of lamin B1 is important for retention of nuclear chromatin during neuronal migration. Proc Natl Acad Sci U S A 110, E1923-1932 (2013).

38. Maske, C.P. et al. A carboxyl-terminal interaction of lamin B1 is dependent on the CAAX endoprotease Rce1 and carboxymethylation. J Cell Biol 162, 1223–1232 (2003).

39. Davies, B.S., Fong, L.G., Yang, S.H., Coffinier, C. & Young, S.G. The posttranslational processing of prelamin A and disease. Annu Rev Genomics Hum Genet 10, 153–174 (2009).

40. Fontecave, M., Atta, M. & Mulliez, E. S-adenosylmethionine: nothing goes to waste. Trends Biochem Sci 29, 243–249 (2004).

41. Luka, Z., Mudd, S.H. & Wagner, C. Glycine N-methyltransferase and regulation of S-adenosylmethionine levels. J Biol Chem 284, 22507–22511 (2009).

42. Dai, Z., Mentch, S.J., Gao, X., Nichenametla, S.N. & Locasale, J.W. Methionine metabolism influences genomic architecture and gene expression through H3K4me3 peak width. Nat Commun 9, 1955 (2018).

43. Mentch, S.J. et al. Histone Methylation Dynamics and Gene Regulation Occur through the Sensing of One-Carbon Metabolism. Cell Metab 22, 861–873 (2015).

44. Cerboni, S. et al. Intrinsic antiproliferative activity of the innate sensor STING in T lymphocytes. J Exp Med 214, 1769–1785 (2017).

45. Prozorovski, T. et al. Sirt1 contributes critically to the redox-dependent fate of neural progenitors. Nat Cell Biol 10, 385–394 (2008).

46. Obata, F. & Miura, M. Enhancing S-adenosyl-methionine catabolism extends Drosophila lifespan. Nat Commun 6, 8332 (2015).

47. Carrasco, M. et al. Glycine N-methyltransferase expression in the hippocampus and its role in neurogenesis and cognitive performance. Hippocampus 24, 840–852 (2014).

48. Obata, F. et al. Nutritional Control of Stem Cell Division through SAdenosylmethionine in Drosophila Intestine. Dev Cell 44, 741–751.e743 (2018).

49. Brady, G.F., Kwan, R., Bragazzi Cunha, J., Elenbaas, J.S. & Omary, M.B. Lamins and Lamin-Associated Proteins in Gastrointestinal Health and Disease. Gastroenterology 154, 1602–1619.e1601 (2018).

50. Kirby, T.J. & Lammerding, J. Emerging views of the nucleus as a cellular mechanosensor. Nat Cell Biol 20, 373–381 (2018).

51. Cho, S. et al. Mechanosensing by the Lamina Protects against Nuclear Rupture, DNA Damage, and Cell-Cycle Arrest. Dev Cell 49, 920–935.e925 (2019).

52. Härtlova, A. et al. DNA damage primes the type I interferon system via the cytosolic DNA sensor STING to promote anti-microbial innate immunity. Immunity 42, 332–343 (2015).

53. Erdal, E., Haider, S., Rehwinkel, J., Harris, A.L. & McHugh, P.J. A prosurvival DNA damage-induced cytoplasmic interferon response is mediated by end resection factors and is limited by Trex1. Genes Dev 31, 353–369 (2017).

54. Yu, Q. et al. DNA-damage-induced type I interferon promotes senescence and inhibits stem cell function. Cell Rep 11, 785–797 (2015).

55. Eguchi, H., Fujiwara, N., Sakiyama, H., Yoshihara, D. & Suzuki, K. Hydrogen peroxide enhances LPS-induced nitric oxide production via the expression of interferon beta in BV-2 microglial cells. Neurosci Lett 494, 29–33 (2011).

56. Zheng, L.S. et al. Mechanisms for interferon-α-induced depression and neural stem cell dysfunction. Stem Cell Reports 3, 73–84 (2014).

57. Miller, R.A. et al. Methionine-deficient diet extends mouse lifespan, slows immune and lens aging, alters glucose, T4, IGF-I and insulin levels, and increases hepatocyte MIF levels and stress resistance. Aging Cell 4, 119–125 (2005).

58. Bruggeman, S.W. et al. Ink4a and Arf differentially affect cell proliferation and neural stem cell self-renewal in Bmi1-deficient mice. Genes Dev 19, 1438–1443 (2005).

59. Sanjana, N.E., Shalem, O. & Zhang, F. Improved vectors and genome-wide libraries for CRISPR screening. Nat Methods 11, 783–784 (2014).

60. Gutscher, M. et al. Real-time imaging of the intracellular glutathione redox potential. Nat Methods 5, 553–559 (2008).

61. Dobin, A. et al. STAR: ultrafast universal RNA-seq aligner. Bioinformatics 29, 15–21 (2013).

62. Liao, Y., Smyth, G.K. & Shi, W. featureCounts: an efficient general purpose program for assigning sequence reads to genomic features. Bioinformatics 30, 923–930 (2014).

63. Love, M.I., Huber, W. & Anders, S. Moderated estimation of fold change and dispersion for RNA-seq data with DESeq2. Genome Biol 15, 550 (2014).

