## Extended Figure 1-6 for "Cellular Stress Signaling Activates Type-I IFN Response Through FOXO3-regulated Lamin Posttranslational Modification"

Extended Data Fig 1.

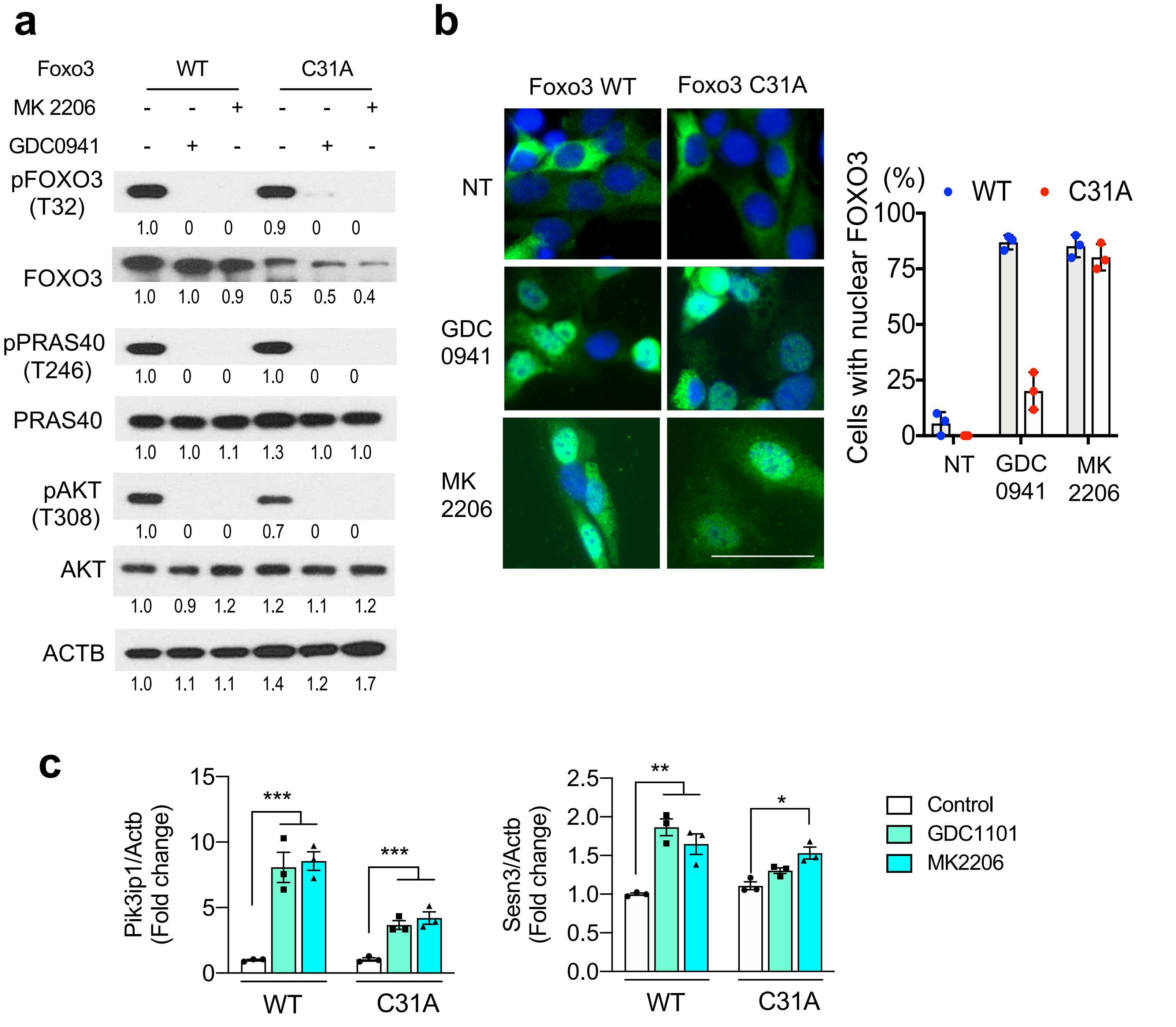


**Extended Data Figure 1. FOXO3 C31A doesn't affect FOXO3 nuclear localization under Akt inhibition. a**. WB analysis of Foxo3 WT or C31A expressing NSPCs. MK2206 (2 μM) or GDC0941 (500 nM) were treated for 0.5 hr. Experiment was repeated three times with similar results. **b**. Microscopic analysis of Foxo3-EGFP following the same treatment as (A). % of cells with nuclear FOXO3 is plotted on the right (n=3). **c**. qRT-PCR analysis for transcriptional targets of FOXO3. Statistical significance was determined by two-way ANOVA for **b** and **c**. Mean ± s.e.m. of 3 independent experiments.***p<0.001, **p<0.01, *p<0.05.

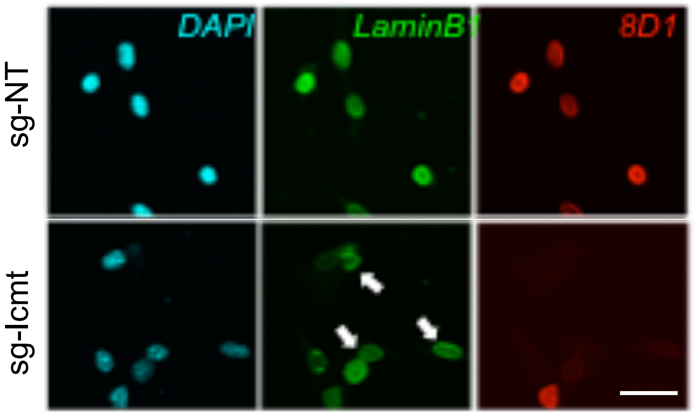


**Extended Data Figure 2. Ablation of ICMT defects prelamin B1 processing.** IF analysis for ICMT depleted NSPC (sg-Icmt) cultures in comparison to control NSPC (sg-NT). Arrows point to nuclei with lamin B1 expression but lacking 8D1 immunoreactivity. Scale bar= 20 μm. Experiment was repeated three times with similar results.

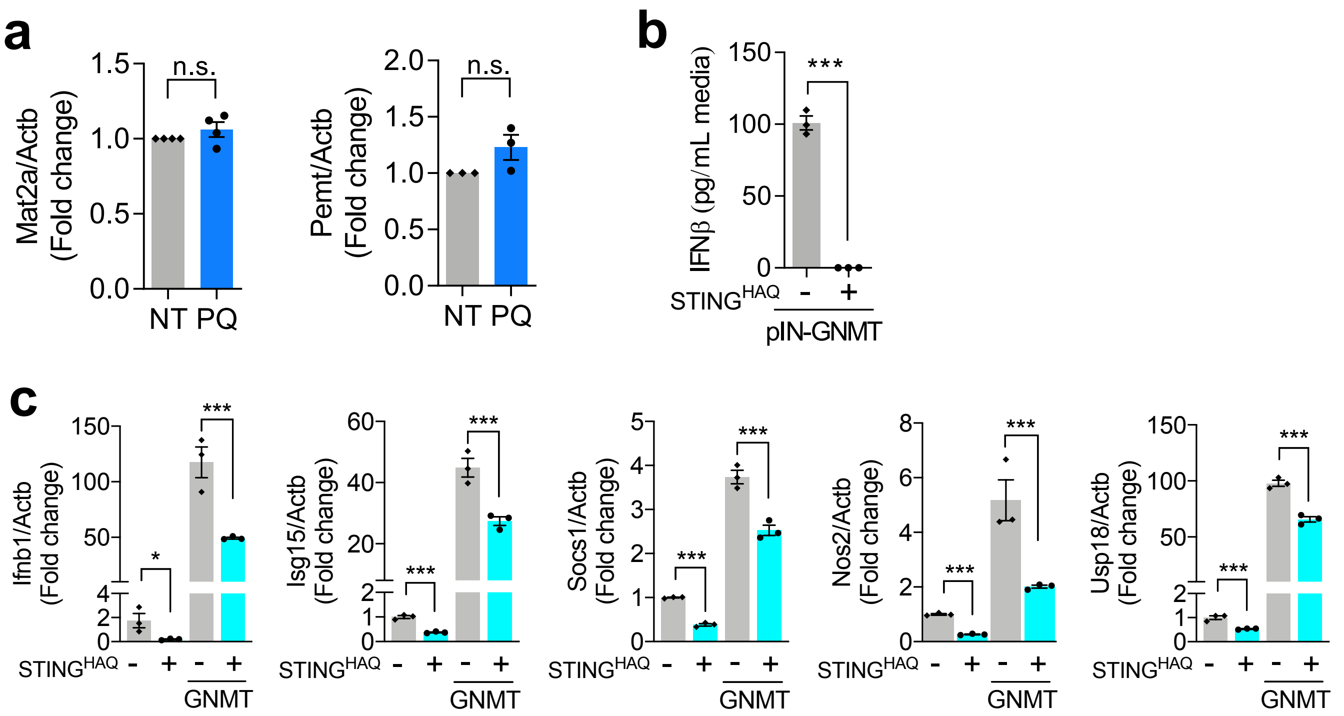


**Extended Data Figure 3. Stress-induced GNMT activates IFN-I response. a.** qRT-PCR results from NSPC stimulated with PQ for 48 hr. Mean ± s.e.m. of 4 independent experiments. **b.** IFNβ secretion in the media following 48 h GNMT induction in $STING^{HAQ}$ mutant expressing NSPCs. Mean ± s.e.m. of 3 independent experiments. **c.** qRT-PCR results from $STING^{HAQ}$ mutant expressing NSPCs for ISGs at 4 days after induction of GNMT. Data are presented the fold change to each control. Mean ± s.e.m. of 3 independent experiments. Statistical significance was determined by unpaired t-test for a and b and by one-way ANOVA for c. ***P<0.001; n.s.= not significant.

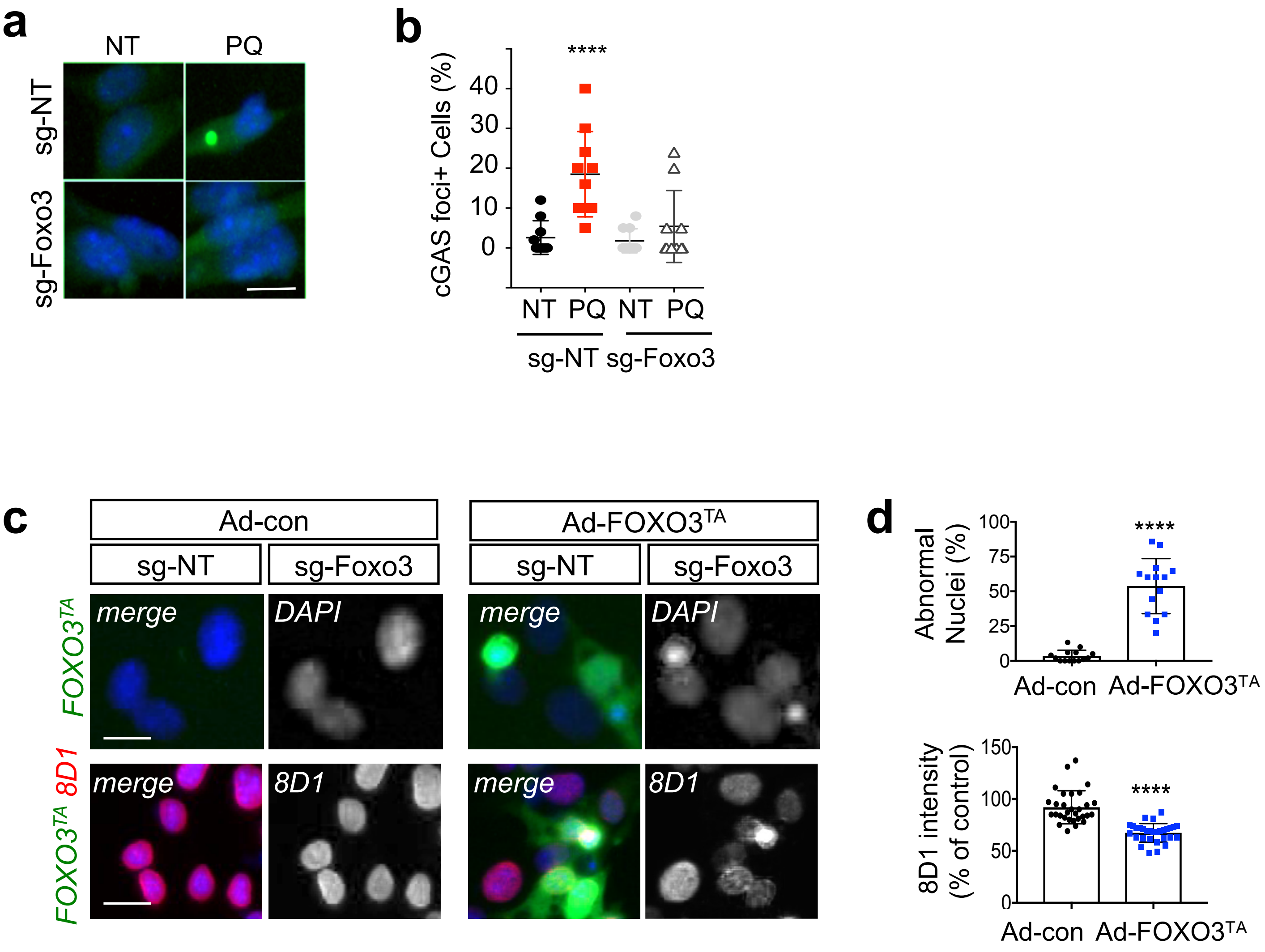


**Extended Data Figure 4. Stress induced-FOXO3 activates IFN-I response and defects lamin processing.** **a.** Microscopic analysis for cGAS-GFP reporter following 1 day of PQ treatment. sg-NT: non-targeted guide RNA, sg-Foxo3: Foxo3 targeted guide RNA. Scale bar= 4 μm. **b**. Quantitation of the percent of cells with cGAS-GFP foci in **a** images. Mean ± s.e.m. of 10 images. **c.** IF analysis for lamin processing (8D1) on either adenovirus for control or FOXO3$^{TA}$ infected NSPC. **d.** Quantitation of the percent of cells with abnormal nuclei and mature lamin B1 (8D1) positive nuclei. Mean ± s.e.m. of 20 images. Statistical significance was determined  y one-way ANPVA for b and by unpaired t-test for d. ****p<0.0001.

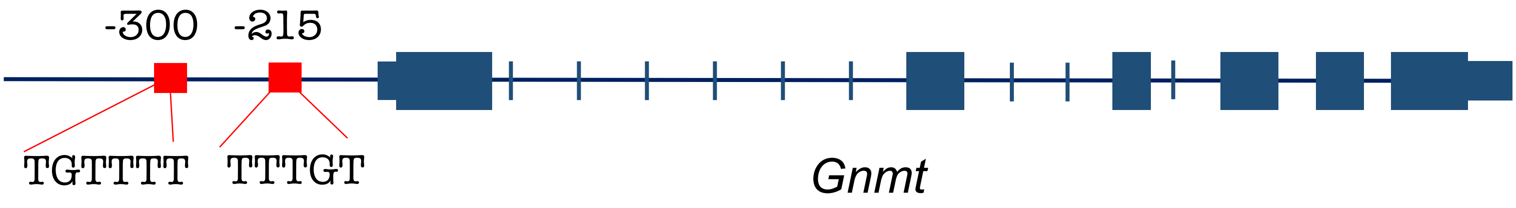


**Extended Data Figure 5. GNMT has FOXO3 binding site on promoter region.** The diagram represents two putative FOXO DBE on mouse GNMT locus.

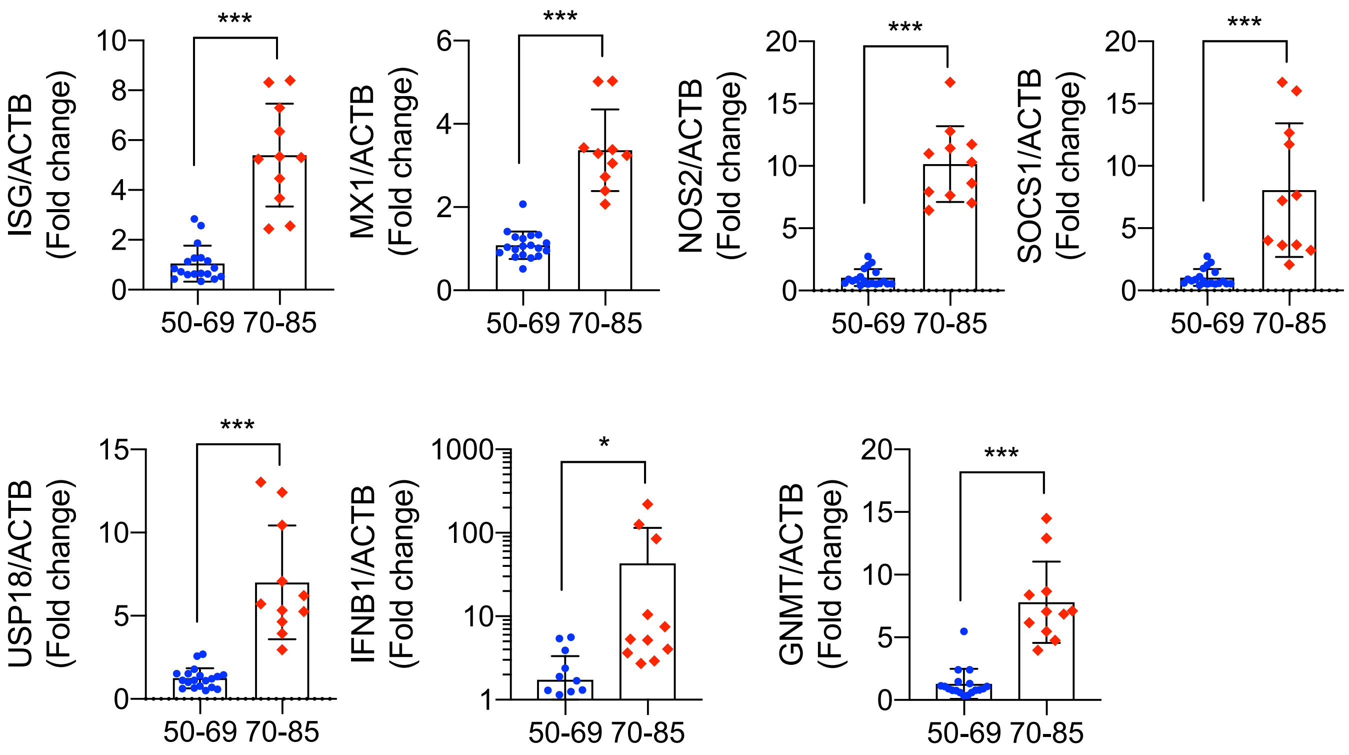


**Extended Data Figure 6. Activated IFN-I response in human aged brains.** qRT-PCR measurement of ISGs and GNMT in cerebellum samples from aging human cohort. Mean ± s.e.m of 11 human brain samples. *P < 0.05; ***P < 0.001. Statistical significance was determined by unpaired t-test.
